# A p97-PML NBs axis is essential for chromatin-bound cGAS untethering and degradation upon senescence-prone DNA damage

**DOI:** 10.1101/2025.10.24.684316

**Authors:** Florent Bressac, Tristan Bargain, Constance Kleijwegt, Wilhelm Bouchereau, Lucas Bonnefoy, Karine Monier, Patrick Lomonte, Armelle Corpet

## Abstract

Cyclic GMP-AMP synthase (cGAS), initially identified as a cytosolic sensor for double-stranded DNA, is now widely recognized as a nuclear protein with distinct STING-independent functions. Its presence in the same compartment as genomic DNA highlights the critical need to regulate its nuclear levels to balance the risk of cell-intrinsic immune activation with the recognition of pathogenic DNA. The recent discovery of a proteasome-dependent degradation mechanism for chromatin-bound cGAS offers new insights into the regulation of nuclear cGAS. In this study, we examine the dynamics and stability of nuclear cGAS following DNA damage. We demonstrate that cGAS is released from chromatin in a process dependent on the p97 segregase, followed by its degradation. When protein degradation is blocked, cGAS accumulates in foci juxtaposed to PML nuclear bodies (PML NBs). We show that this juxtaposition is SUMO-dependent, with both SUMO and PML required for cGAS degradation, a process that also involves the Cullin3-RING E3 ubiquitin Ligase complex. Increasing cGAS levels on chromatin after damage by preventing its degradation expands its localization on chromatin, correlating with a dampened DNA damage response and impaired senescence entry. Overall, our findings show that a p97-PML NBs axis modulates cGAS abundance, ensuring proper regulation of the DNA damage response and balancing the possible cell-intrinsic activation of innate immune responses with senescence entry.

## Introduction

The innate immune response is the first line of defense against infections. Cells have evolved a number of specialized immunity receptors called PRRs (Pattern Recognition Receptors) that recognize pathogen-specific patterns and induce intracellular signaling pathways leading to an inflammatory response to fight pathogen infection ^1^. Among the various PRRs, the cyclic GMP-AMP synthase (cGAS, also known as MB21D1), was identified as an essential cytosolic DNA sensor which plays a key role in response to microbial infections ^2^. Upon binding to cytosolic double-stranded DNA such as foreign viral DNA, cGAS undergoes a conformational change and dimerizes to catalyze the synthesis of cGAMP, a cyclic dinucleotide that induces interferon type I (IFN-I) and innate immunity responses through a STING-TBK1-IRF3 axis ^3^. Due to its inherent property to recognize DNA in a sequence-independent manner, cGAS can also be activated by DNA that has been mislocalized following genotoxic stresses. This self-DNA, leaking from the nucleus in the form of micronuclei ^4,5^, cytosolic chromatin fragments (CCFs) ^6,7^ or from mitochondrial sources ^8,9^, can potentially trigger aberrant activation of cGAS, which contributes to autoimmunity, inflammation as well as senescence. It is thus essential for cells to maintain a delicate balance between detecting pathogenic DNA and keeping cGAS inactive in response to host DNA.

It is now well established that a great proportion of cGAS is nuclear ^10,11^. Structural studies have revealed that the tight tethering of cGAS to the acidic patch of the histones, the main protein component of the nucleosome, is key to prevents its auto-reactivity towards self-DNA ^12–17^. In addition, a growing number of studies have uncovered various STING-independent functions for cGAS in the nucleus such as inhibition of DNA repair ^18,19^, regulation of DNA replication ^20^ or nuclear envelope repair ^21^. Remarkably, nuclear cGAS is maintained at low levels during interphase by a specific degradation mechanism ^22^. The Cullin-RING E3 ubiquitin ligase 5 complex (CRL5), and its substrate receptor SPSB3, are essential mediators for the proteasome-dependent degradation of nucleosome-bound cGAS ^22^.

Upon DNA damage, chromatin architecture is dramatically altered. While chromatin unfolding/decompaction contributes to the access and timely repair of DNA lesions, histone dynamics also participates in the restoration of epigenome integrity ^23^. Chromatin-bound cGAS inhibits double-strand breaks (DSBs) repair by homologous recombination, notably by preventing the recruitment of RAD51 through DNA compaction ^18,19^. Binding of the MRE11-RAD50-NBS1 complex to DNA damage sites has a crucial role in releasing cGAS from nucleosome sequestration to enable its activation by dsDNA in the cytoplasm ^24^. However, the mechanisms underlying the regulation of nuclear cGAS activity and its nuclear functions are still poorly defined. In particular, how cells manage nucleoplasmic cGAS released from chromatin after DNA damage and whether this could influence the early onset of senescence, remains unexplored.

Within the densely packed nuclear space, PML Nuclear Bodies (PML NBs), form membrane-less organelles (MLOs) of 0.1-1μm diameter which, by concentrating proteins at specific places, play pivotal roles in controlling biochemical reactions in space and time ^25,26^. PML NBs have been both implicated in the regulation of essential physiological responses such as DNA repair and senescence entry, and in protein-degradation mechanisms ^27^, thus placing them as ideal candidates to maintain nuclear protein homeostasis upon DNA damage. One component of PML NBs ^28^, the ubiquitin-dependent unfoldase/segregase p97, also known as VCP, is a central component of the ubiquitin-proteasome system. With the help of cofactors, p97 binds to ubiquitinated substrates and, using its ATPase activity, processes them by unfolding or extracting them from various cellular locations, including chromatin ^29^. Processed proteins can then be addressed to the proteasome for their degradation illustrating the essential function of p97 in regulating protein homeostasis.

Here, we investigate the dynamics and stability of nuclear cGAS upon DNA damage in human primary fibroblasts. Upon DSB induction, we show that cGAS is untethered from chromatin and degraded in a proteasome-dependent manner. Preventing cGAS degradation triggers its accumulation in foci that juxtapose PML NBs and proteasome. The p97 segregase is essentiel for cGAS release from chromatin and foci formation next to PML NBs, which are then involved in cGAS degradation together with a Cullin3-RING E3 ubiquitin Ligase complex. While cGAS localizes mainly on specific heterochromatin regions, inhibition of its release from chromatin correlates with its spreading on euchromatin and a dampened DNA damage response, impairing senescence entry. We thus unveil a novel p97-PML NBs axis that controls the nucleoplasmic pool of cGAS upon DNA damage to fine-tune the risk of autoimmunity activation and senescence entry.

## Results

### Chromatin destabilization mobilizes cGAS from chromatin and leads to its degradation

cGAS is constitutively inactivated by its tight binding to nuclear chromatin which prevents its binding to dsDNA and dimerization ^12–17^. While DNA damage triggers its MRN-dependent release from chromatin to activate a cytosolic dsDNA sensing and inflammatory response ^24^, what happens to the pool of remaining nuclear cGAS after DNA damage is currently unknown. As a first steps towards deciphering the dynamics of nuclear cGAS after DNA damage, we examined its levels in human primary foreskin diploid fibroblasts BJ cells treated with etoposide, a well-established inhibitor of topoisomerase II which triggers DNA damage in the form of double-strand breaks (DSBs) ^30^. The use of primary cells is key to investigate the physiological response to DNA damage which can ultimately lead to senescence entry, a pathway that is inactivated in cancerous cells ^31^. Treatment of BJ cells with etoposide for 24 hours triggers, as expected, a strong accumulation of γH2AX, a marker of DSBs (Figures 1A-B). In parallel, a strong decrease in the nuclear signal of cGAS was observed by immunofluorescence (Figure 1A), concomitant with a significant decrease in the total pool of cGAS by western blotting (Figure 1B), without diminishing cGAS mRNA levels (Sup. Figure 1A). Use of leptomycin B, an inhibitor of the nuclear export machinery, did not rescue cGAS levels in the nucleus (Sup. Figure 1B), suggesting a potential nuclear degradation of cGAS upon DNA damage. In order to further substantiate our observations, we treated BJ cells with TSA, a well-known HDAC inhibitor which triggers the opening of chromatin condensed regions ^32^, on which cGAS is particularly bound ^33^. TSA induced a strong decrease of cGAS presence in the nucleus as well as a significant reduction in its total amount in the cells, without any significant accumulation of γH2AX (Figures 1A-B), thus extending our findings to other chromatin destabilizing agents beyond DNA damage. To track cGAS localization more thoroughly, we established cells stably expressing an epitope-tagged version of cGAS by lentiviral transduction. While cGAS levels are low in primary BJ cells, expression of higher levels of GFP-cGAS could be achieved by doxycyclin induction (Sup. Figure 1C). Cellular fractionation confirmed the tight tethering of GFP-cGAS to the chromatin (Figure 1C) and its low abundance in the cytosolic and nuclear soluble fractions in non-treated cells. Addition of etoposide or TSA induced a significant decrease of the chromatin-bound pool of cGAS, without any major increase in the other fractions (Figure 1C). cGAS mobilization from chromatin following DNA damage was confirmed on the endogenous cGAS (Figure 1D). In contrast, we could not detect any significant effect of etoposide on cGAS nuclear localization/levels in the HeLa cancerous cell line (Sup. Figures 1D-F). Thus, our data suggest that, in primary cells competent for senescence induction, chromatin destabilization and opening release a fraction of chromatin-bound cGAS, which is subsequently degraded.

**Figure 1:**
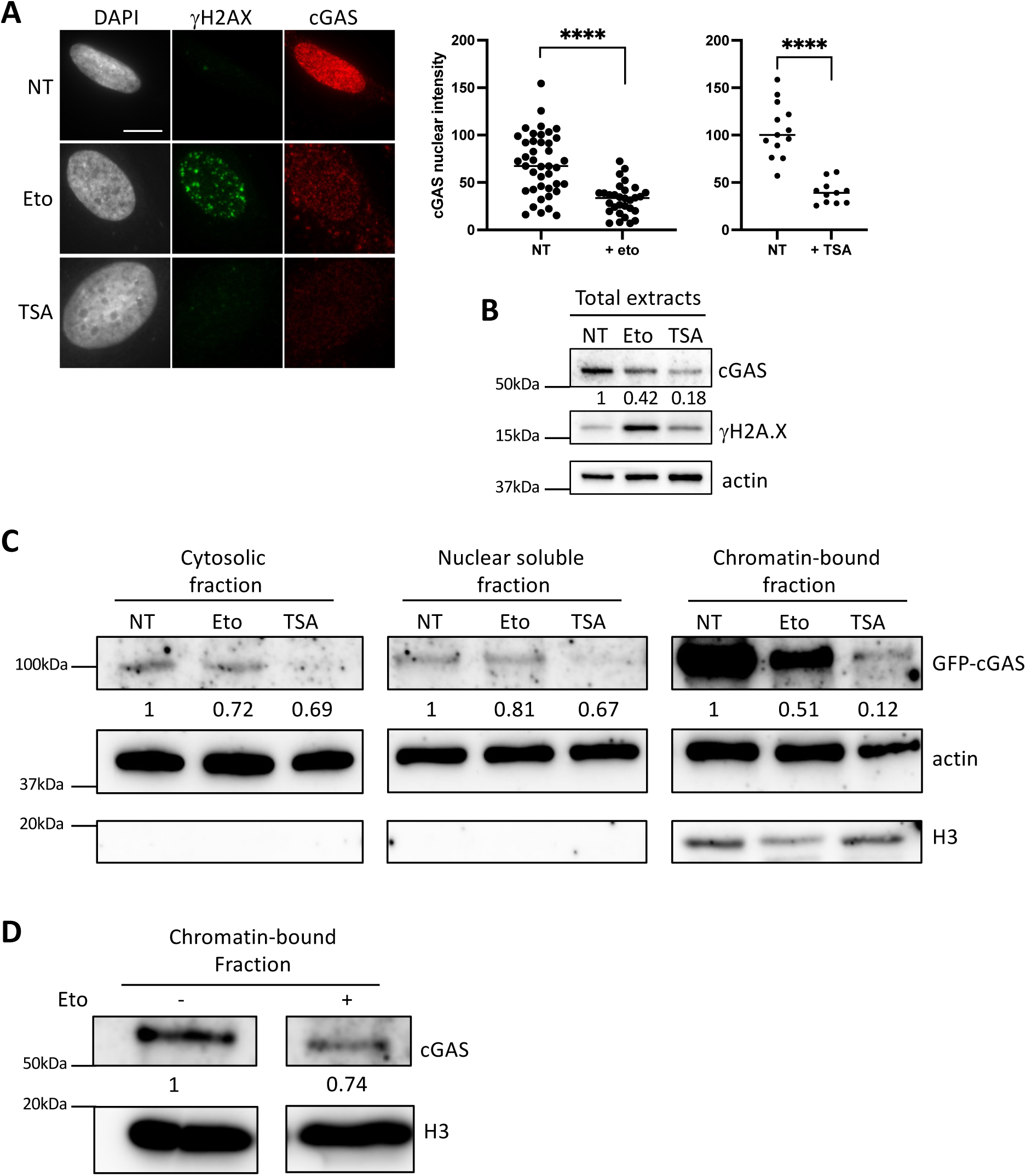
Chromatin destabilization induces a global and chromatin-bound decrease of cGAS levels in primary cells. (A) (left) Fluorescence microscopy visualization of cGAS (red) and γH2AX (green) in BJ cells treated with etoposide at 10μM for 24h or TSA at 2uM for 24h. Cell nuclei are visualized by DAPI staining (grey). Scale bar is 10μm. (Right) Plot shows quantitative analysis of nuclear cGAS intensity in cells. P-values (Student t-test): ****<0,0001. (B) Western-blot visualization of cGAS from total cell extracts on BJ cells treated with the same conditions. Actin is a loading control and quantification of cGAS levels relative to actine are shown below the WB (numbers are representative from three independent experiments). (C) Western-blot visualization of cGAS from different fractions of BJ GFP-cGAS WT cells treated with etoposide at 10μM for 24h or TSA at 2μM for 24h and Doxycycline at 100ng/mL for 24h. H3 is a loading control for the chromatin-bound fraction. Quantification of cGAS levels relative to actin levels (cytosolic and nuclear fractions) or relative to H3 (chromatin-bound fraction) are shown below. (D) Western-blot visualization of cGAS and H3 in the chromatin-bound fraction of BJ cells treated with etoposide at 10μM for 24h. Quantification of cGAS levels relative to H3 are shown below.

### Proteasome inhibition leads to the accumulation of cGAS in foci juxtaposed to PML NBs

While cGAS degradation mechanisms have extensively been studied in the cytosol ^34^, its turn-over in the nucleus remains under-investigated. The recent characterization of the CRL5-SPSB3 complex required for the proteasome-dependent degradation of chromatin-bound cGAS in a cell-cycle regulated manner in non-stressed cells ^22^, has opened up new perspectives on the importance of regulating cGAS nuclear levels. Addition of MG132, a proteasome inhibitor, in the last 8 hours of etoposide treatment, restored the total and chromatin-bound pool of cGAS after damage (Figures 2A-B) and increased cGAS pan-nuclear levels as visualized by immunofluorescence of cGAS (Figure 2C). Remarkably, nuclear cGAS formed intense foci in the nucleus, that juxtaposed perfectly next to PML NBs (Figure 2C). Addition of MG132 following TSA treatment was sufficient to restore cGAS nuclear/total levels, and also led to the appearance of cGAS foci juxtaposed to PML NBs (Sup. Figures 2A-B). While all PML NBs were not juxtaposed to cGAS foci, on the contrary, every single cGAS spot was juxtaposed to a PML NB. Quantification of the distance between cGAS and PML signal revealed a minimal distance well below 200nm underscoring the close spatial proximity of cGAS foci and PML NBs (Figure 2C).

**Figure 2:**
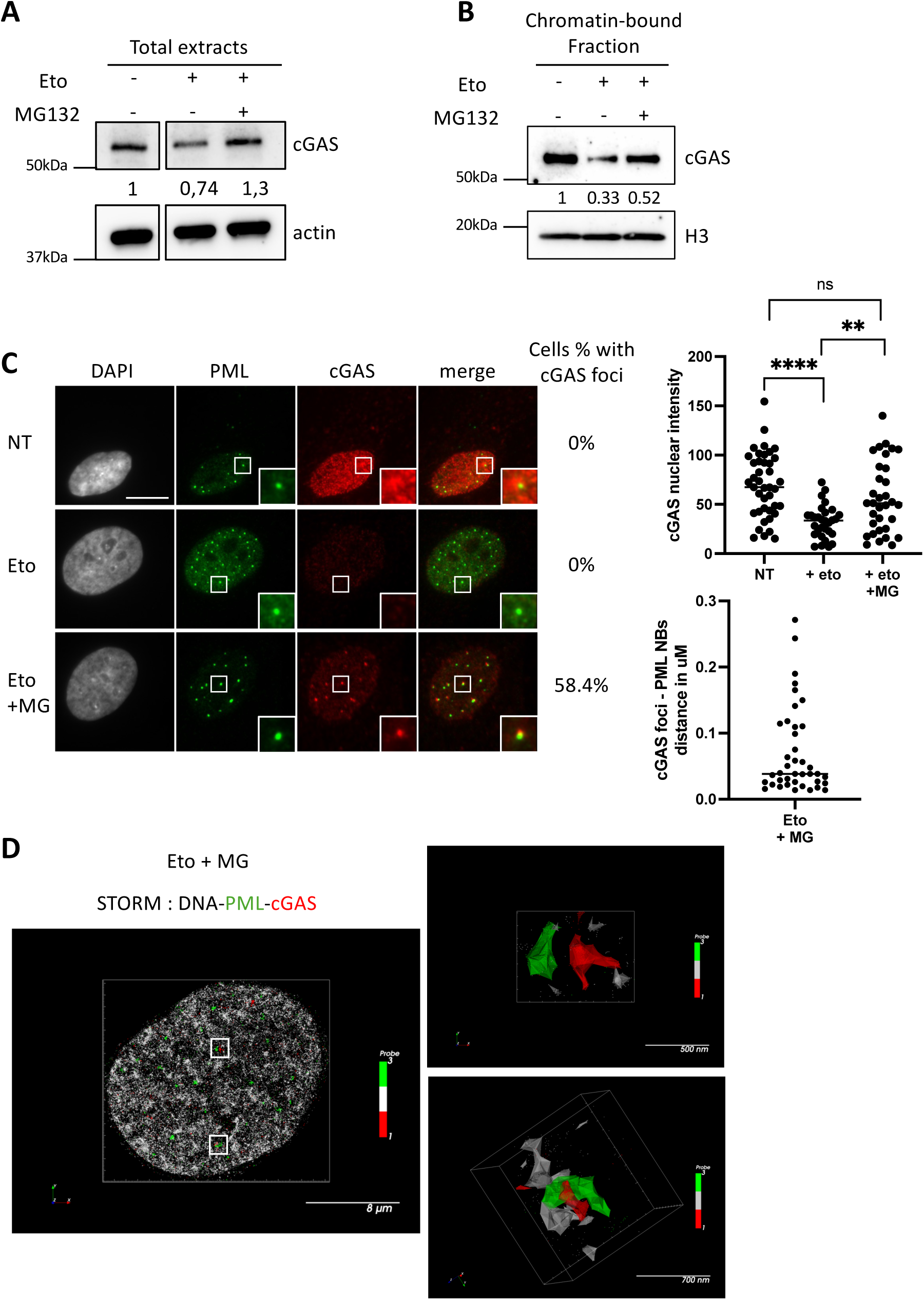
Proteasome inhibition prevents the decrease of cGAS in the nucleus and leads to the formation of cGAS foci juxtaposed to PML NBs. (A) Western-blot visualization of cGAS from total cell extracts on BJ cells treated with etoposide at 10μM for 24h and MG132 at 2,5μM for the last 8h. Actin is a loading control and quantification of cGAS levels relative to actin are shown below the WB (numbers are representative from three independent experiments). (B) Western-blot visualization of cGAS from the chromatin-bound fraction of BJ cells treated as in (A). H3 is a loading control and quantification of cGAS levels relative to H3 are shown below. (C) (left) Fluorescence microscopy visualization of PML (green) and cGAS (red) in BJ cells treated as in (A). Cell nuclei are visualized by DAPI staining (grey). Proportion of cells showing cGAS juxtaposed to PML NBs is shown on the right (numbers are representative from 3 independent experiments). Scale bar is 10μm. Insets represent enlarged images (3X) of selected areas. (Right) Upper plot shows quantitative analysis of nuclear cGAS intensity from immunofluorescence images. P-values (Student t-test) : **<0,01 ; ****<0,0001, ns : non-significant. Lower plot shows distance in μm between the center of cGAS foci and the center of the closest PML NB. (D) Super-resolution microscopy image by dSTORM showing DNA (gray), PML (green) and cGAS (red) as point by point vizualisation in a single nucleus from BJ cells treated with etoposide at 10μM for 24h and MG132 at 2,5μM for the last 8h. Enlarged insets are visualized as 3D-cluster hulls on the right. Scale bars are indicated below each image.

cGAS foci observed upon proteasome inhibition may correspond either to undegraded nucleoplasmic cGAS or to a retained chromatin-bound fraction. Given its known binding to centromeric satellite DNA ^33^, we first examined whether non-degraded nuclear cGAS foci may colocalize with centromeres. Use of the anti-centromeres antibodies human ACA unequivocally showed that cGAS foci do not accumulate on/close to centromeres (Sup. Figure 2C). To further substantiate the subnuclear localization of cGAS foci, we performed super-resolution imaging using dSTORM (direct Stochastic Optical Reconstruction Microscopy) in cells treated with etoposide and MG132 and labelled with the highly specific DNA probe SPY555-DNA, which colocalizes with H2A-H2B-GFP (Sup. Figure 2D). Three-color dSTORM images of cGAS and PML with DNA showed that cGAS foci do not colocalize with DNA, but are instead positioned in close proximity to PML NBs (Figure 2D), as observed by standard microscopy (Figure 2C). These observations suggest that cGAS foci correspond to nucleosoluble cGAS sequestered near PML NBs prior to degradation. While PML NBs have been shown to juxtapose DNA damage foci ^35^, their close proximity with γH2AX was exclusive with cGAS foci (Sup. Figure 2E). Thus, we conclude that the ubiquitin-proteasome system (UPS) is essential to regulate the amount of nucleoplasmic cGAS following stresses destabilizing the chromatin structure. cGAS accumulation in foci juxtaposed to PML NBs upon proteasome inhibition suggests an implication of these membrane-less organelles in the degradation process.

### PML NBs are implicated in cGAS degradation but not foci formation

PML NBs have been implicated in protein degradation, particularly through their interaction with components of the UPS. Upon arsenic-induced stress, which greatly enhances PML SUMOylation, ubiquitinated proteins as well as subunits of the proteasome localize within PML NBs, which sustains catabolism of SUMOylated proteins ^36^. Using the FK2 antibody which recognizes mono- and poly-ubiquitinated proteins, we first verified that ubiquitinated proteins accumulate in PML NBs upon addition of MG132 following etoposide treatment (Sup. Figure 3A). cGAS foci closely juxtapose FK2 foci (Sup. Figure 3B) as expected given their juxtaposition to PML NBs (Figure 2C). We then examined the localization of the proteasome subunit, ADRM1 (the human homolog of yeast Rpn13), a component of the 19S regulatory particle (RP) responsible for Ub recognition and unfolding substrates, which is juxtaposed to PML NBs upon replicative senescence ^37^. In untreated cells, ADRM1 forms a few foci in the nucleus, that juxtapose neither PML NBs nor cGAS (Sup. Figure 3C). Upon DNA damage induction and proteasome inhibition, 59.4% +/- 10.4% of ADRM1 foci juxtaposed to PML NBs. Moreover, triple immunofluorescence staining revealed that cGAS foci closely associate with PML-ADRM1 foci (Figure 3A), suggesting a key role of PML NBs as platforms for the proteasomal degradation of cGAS.

**Figure 3:**
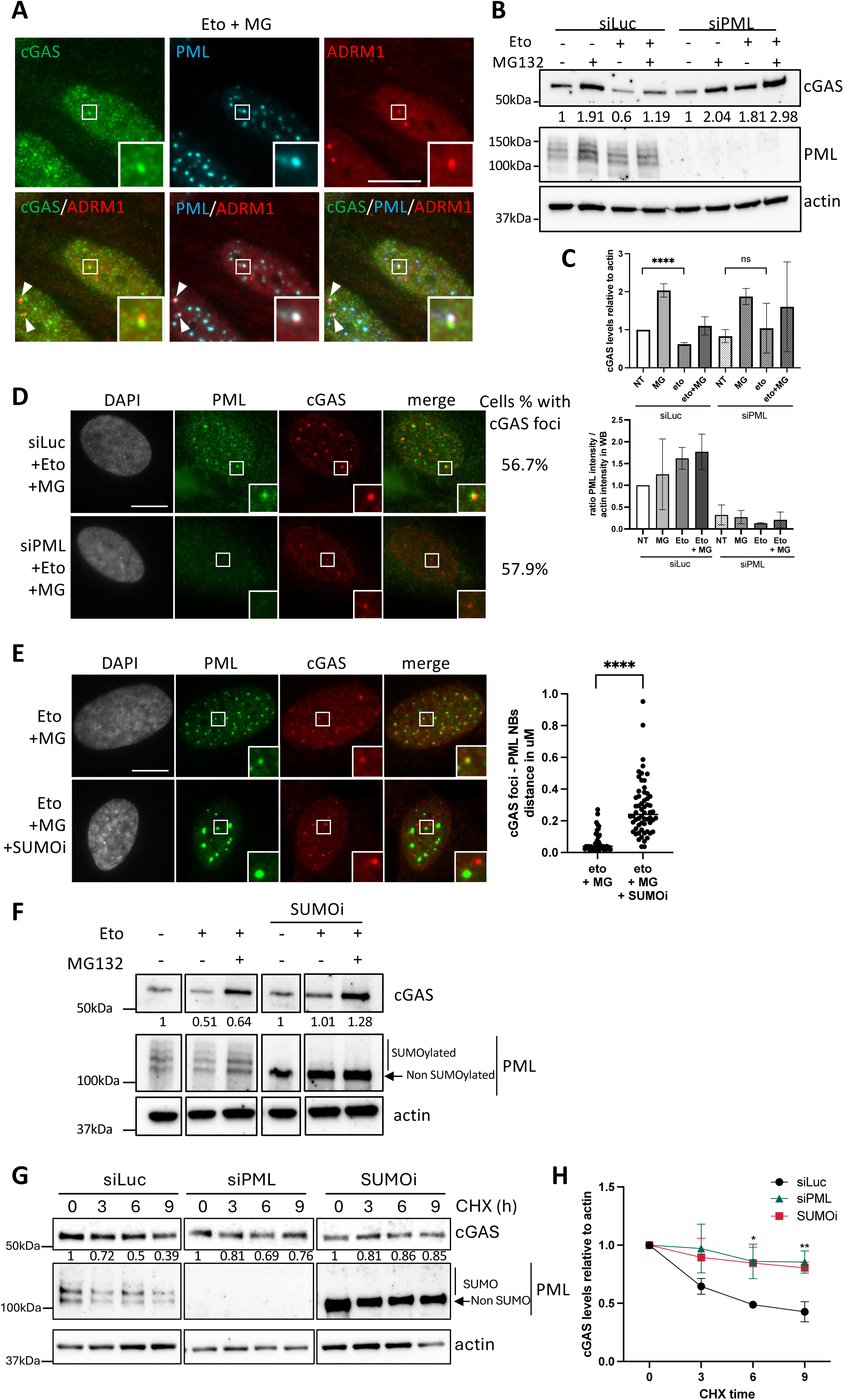
PML NBs are implicated in cGAS degradation but not foci formation. (A) Fluorescence microscopy visualization of cGAS (green), PML (cyan) and ADRM1 (red) in BJ cells treated with etoposide at 10μM for 24h and MG132 at 2,5μM for the last 8h. Insets represent enlarged images (3X) of selected areas showing cGAS-ADRM1 and PML-ADRM1 juxtaposition. Arrowheads also indicate mark these juxtapositions. Scale bar is 10μm. (B) Western-blot visualization of cGAS and PML from total cell extracts on BJ cells treated with etoposide at 10μM for 24h, MG132 at 2,5μM for 8h and siLuc or siPML for 48h. Actin is a loading control and quantification of cGAS levels relative to actin is shown below this representative WB. (C) The two histograms show quantification of cGAS and PML levels from three independent experiments (cGAS) or two independent experiments (PML). P-values (Student t-test) : ****<0,0001, ns : non-significant. (D) Fluorescence microscopy visualization of PML (green) and cGAS (red) in BJ cells treated as in (B). Cell nuclei are visualized by DAPI staining (grey). Scale bar is 10μm. Insets represent enlarged images (3X) of selected areas. (E) (Left) Fluorescence microscopy visualization of PML (green) and cGAS (red) in BJ cells treated with etoposide at 10μM for 24h, SUMOi (ML-792, a SUMO-Activating Enzyme inhibitor) at 1μM for 8h and with MG132 at 2,5μM for 8h. Cell nuclei are visualized by DAPI staining (grey). Scale bar is 10μm. Insets represent enlarged images (3X) of selected areas. (Right) Plot shows quantification of the distance in μm between the center of cGAS foci and the center of the closest PML NB. P-values (Student t-test): ****<0,0001. (F) Western-blot visualization of cGAS and PML from total cell extracts on BJ cells treated as in (E). Actin is a loading control and quantification of cGAS and PML levels relative to actin are shown below the WB. (G) Western-blot visualization of cGAS and PML from total cell extracts on BJ cells treated with cycloheximide at 100μg/mL for the indicated hours, with etoposide at 10μM for 24h and siLuc or siPML for 48h or with SUMOi at 1μM for 8h. Actin is a loading control and quantification of cGAS levels relative to actin is shown below this representative WB. (H) Line plot showing quantification of cGAS levels relative to actin from 3 independent experiments as performed in (E). P-values (Student t-test) : * <0,05 ; **<0,01.

To investigate the role of PML NBs in controlling cGAS nuclear levels, PML was depleted by a targeted RNAi-based approach, as in ^38^. Western blot analysis confirmed the significant decrease in PML total levels in siPML-treated cells compared to the control siRNA (Figures 3B-C). While PML knock-down did not significantly altered cGAS total levels in non-treated cells, it abrogated the previously observed decrease of total cGAS upon addition of etoposide. Addition of MG132 in presence of DNA damage led to an even greater restoration of cGAS levels (Figures 3B-C). These data confirm that PML is required to degrade nuclear cGAS following its mobilization from chromatin upon DNA damage. Close examination of siPML-depleted cells revealed that cGAS foci remained present in the nuclei, indicating that their formation is independent of PML (Figure 3D) and may happen before juxtaposition with PML NBs.

Protein localization within/next to PML NBs is intimately linked with SUMOylation. UBC9, the sole SUMO E2-conjugating enzyme localizes withing PML NBs where it SUMOylates proteins which may help in the retention of proteins through binding to the PML SUMO-interacting Motif (SIM) ^26^. To investigate the role of SUMOylation for cGAS nuclear dynamics upon proteasome inhibition, we conducted experiments in the presence of ML-792, a SUMO activating enzyme (SAE) inhibitor, referred to as SUMOi, which prevents protein SUMOylation. SUMOi altered PML NBs structure resulting in the appearance of enlarged PML NBs (Figure 3E and 39), mimicking the shape of PML NBs upon SUMO1/2/3 depletion ^38^. Accordingly, SUMOi prevented PML SUMOylation (Figure 3F). In conditions of proteasome inhibition, cGAS foci formation could still occur in absence of SUMOylation, but their juxtaposition next to PML NBs was lost (Figure 3E), consistent with our previous results showing that cGAS foci can form independently of PML NBs. Mean minimal distance between cGAS foci and PML NBs increased from 0,071 to 0,28μm as shown by quantification (Figure 3E), thus underscoring the loss of juxtaposition. Given that PML NBs are implicated in cGAS degradation, preventing its juxtaposition to PML NBs should alter its degradation. Indeed, addition of SUMOi blunted the decrease of cGAS levels after etoposide treatment (Figure 3F). We next employed cycloheximide in combination with PML-targeting siRNA or the SUMOi to test whether PML or SUMOylation regulate cGAS stability upon etoposide treatment. Consistent with a role in controlling cGAS nucleosoluble levels (Figures 3B, 3F), both PML depletion or SUMO inhibition significantly increased the half-life of cGAS in cycloheximide chase experiments at 6 and 9 hours after treatment (Figures 3G-H). Together, these data show that SUMOylation plays an essential role in targeting/retaining cGAS juxtaposed to PML NBs, which may then participate in cGAS degradation.

### NEDDylation and Cullin3 participate in cGAS degradation following DNA damage

We next aimed to characterize the complex responsible for cGAS degradation by the proteasome. Cullin-RING E3 Ligases (CRLs) are the largest superfamily of E3 Ubiquitin ligases, involved in targeting proteins for degradation. Since all cullins are activated through the covalent attachment of NEDD8, we first used a broad inhibitor of NEDDylation, MLN4024, here abbreviated as NEDD8i, to target the NEDD8 Activating Enzymes (NAE) and thus inhibit CRLs. In cells treated with etoposide, addition of NEDD8i triggered cGAS foci formation juxtaposed to PML NBs, and alleviated the nuclear decrease of cGAS observed upon etoposide treatment alone (Figure 4A). These results are reminiscent of those observed with the proteasome inhibitor MG132 (Figure 2C), suggesting that CRLs are implicated in cGAS degradation. Since cGAS was recently found to be degraded by the SPSB3-CRL5 complex ^22^, we first checked whether this complex impacted on the regulation of cGAS total levels and cGAS foci formation after DNA damage. Removal of SPSB3 or CUL5 by RNAi treatment (Sup. Figure 4A), did not induce cGAS foci formation in non-treated cells (Sup. Figures 4B-C). Upon DNA damage combined with proteasome inhibition, knock-down of SPSB3 or CUL5 had no impact on cGAS foci formation, as shown by quantification (Sup. Figure 4D), suggesting the existence of an alternative specific pathway for nuclear cGAS degradation upon DNA damage. We next undertook a candidate-based approach to investigate whether any other cullin could be implicated in degradation of cGAS. Using published siRNA sequences against Cul1, 2, 3, 4A and 4B, we first verified efficiency of siRNA depletion by RT-QPCR or WB analysis (Figure 4B, Sup. Figures 4A, E). Cul1, 2, 4A, or 4B knock-down did not trigger the formation of cGAS foci juxtaposing PML NBs (Sup. Figure 4B). In contrast, depletion of Cul3, by two independent sets of siRNAs, induced an accumulation of cGAS in foci juxtaposing PML NBs (Figure 4B, Sup. Figures 4B, F), and diminished cGAS decrease upon etoposide treatment (Figure 4C). Thus, our results identify Cul3 as a new important player in cGAS degradation, mediated by a PML-NBs - proteasome axis.

**Figure 4:**
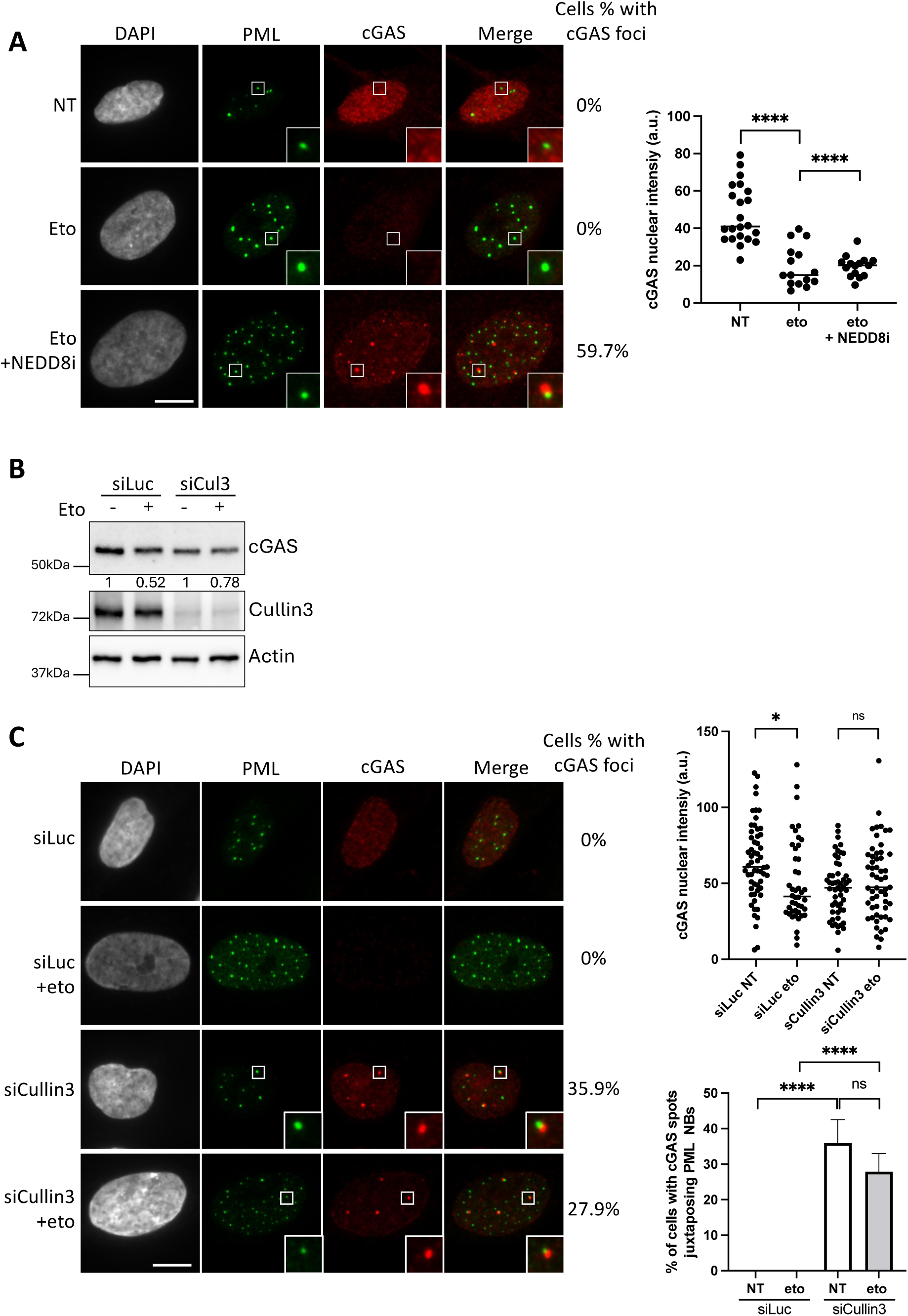
NEDDylation and Cullin3 are implicated in cGAS degradation. (A) Fluorescence microscopy visualization of PML (green) and cGAS (red) in BJ cells treated with etoposide at 10μM for 24h and NEDD8i (MLN4924, a NEDDylation Activating Enzyme inhibitor), at 1μM for 8h. Plot on the right shows quantitative analysis of nuclear cGAS intensity from immunofluorescence images (one representative experiment). P-values (Student t-test): ****<0,0001. (B) Western-blot visualization of cGAS, Cullin3 or Actin from total cell extracts on BJ cells treated with etoposide at 10μM for 24h, MG132 at 2,5μM for 8h and siLuc or siCullin3_01 for 48h. Actin is a loading control and the mean quantification of cGAS levels relative to actin from 3 independent experiments is shown below the WB. (C) (Left) Fluorescence microscopy visualization of PML (green) and cGAS (red) in BJ cells treated as in (B). Cell nuclei are visualized by DAPI staining (grey). Scale bar is 10μm. Insets represent enlarged images (3X) of selected areas. (Right) Upper plot shows quantitative analysis of nuclear cGAS intensity from immunofluorescence images (n=2 independent experiments). P-values (Student t-test) : *<0,05 ; ns : non-significant. Lower plot shows the percentage of cells showing cGAS foci juxtaposing PML NBs (n=3 independent experiments). P-values (Student t-test) : ****<0,0001 ; ns : non-significant.

### p97 is implicated in the release of chromatin-bound cGAS following DNA damage

We next asked whether a protein complex might mediate cGAS release from chromatin following chromatin relaxation upon DNA damage or TSA treatment. The p97 segregase, which is a component of PML NBs ^28^, is recruited to DNA damage sites where it disassembles ubiquitinated protein substrates, ensuring protein turnover and the regulation of chromatin structure around DSBs which in turn, helps to preserve genome stability ^29^. We thus investigated the role of p97 in the regulation of cGAS protein turnover after DNA damage. We first monitored the total pool of cGAS, when p97 ATPase activity was blocked, by acute chemical inhibition with CB5083, here abbreviated as p97i. Inactivation of p97 blunted the decrease of total cGAS levels observed upon etoposide addition in the non-treated control cells (Figure 5A). Addition of MG132 restored even higher cGAS levels after etoposide treatment (Figure 5A), as compared to the cells without p97i (Figure 2A). In immunofluorescence, treatment with p97i abolished the decrease in cGAS nuclear intensity observed in etoposide-treated cells (Figure 5B). Since p97 is required to extract proteins from chromatin after damage, we then looked at cGAS chromatin-bound levels by cellular fractionation. The reduction of cGAS chromatin-bound levels upon etoposide addition was abrogated when p97 is inhibited (Figure 5C), demonstrating a key role for p97 in regulating cGAS chromatin-bound levels after DNA damage. Addition of p97i also abolished the decrease in total cGAS or nuclear cGAS (Sup. Figures 5B-C), and partially restored cGAS chromatin-bound levels upon TSA treatment (Sup. Figure 5D), confirming the importance of p97 in cGAS untethering upon chromatin mobilization. Since cGAS forms nuclear foci upon proteasome inhibition (Figure 2C), independently of PML (Figure 3D) and without colocalization with DNA (Figure 2D), we sought to strengthen the hypothesis that, these foci arise from undegraded nucleoplasmic cGAS after its release from chromatin. We thus inspected cGAS foci formation in absence of p97 activity, which inhibits cGAS untethering from chromatin. In conditions of p97i alone, no foci were observed for cGAS (Sup. Figure 5A). Upon DNA damage and proteasome inhibition, foci formation of cGAS was lost upon p97 inhibition, in contrast to the situation without p97i (Figure 5D). These results suggest that cGAS foci formation occurs downstream of p97 activity and involves the accumulation of untethered, nucleoplasmic cGAS upon proteasome inhibition.

**Figure 5:**
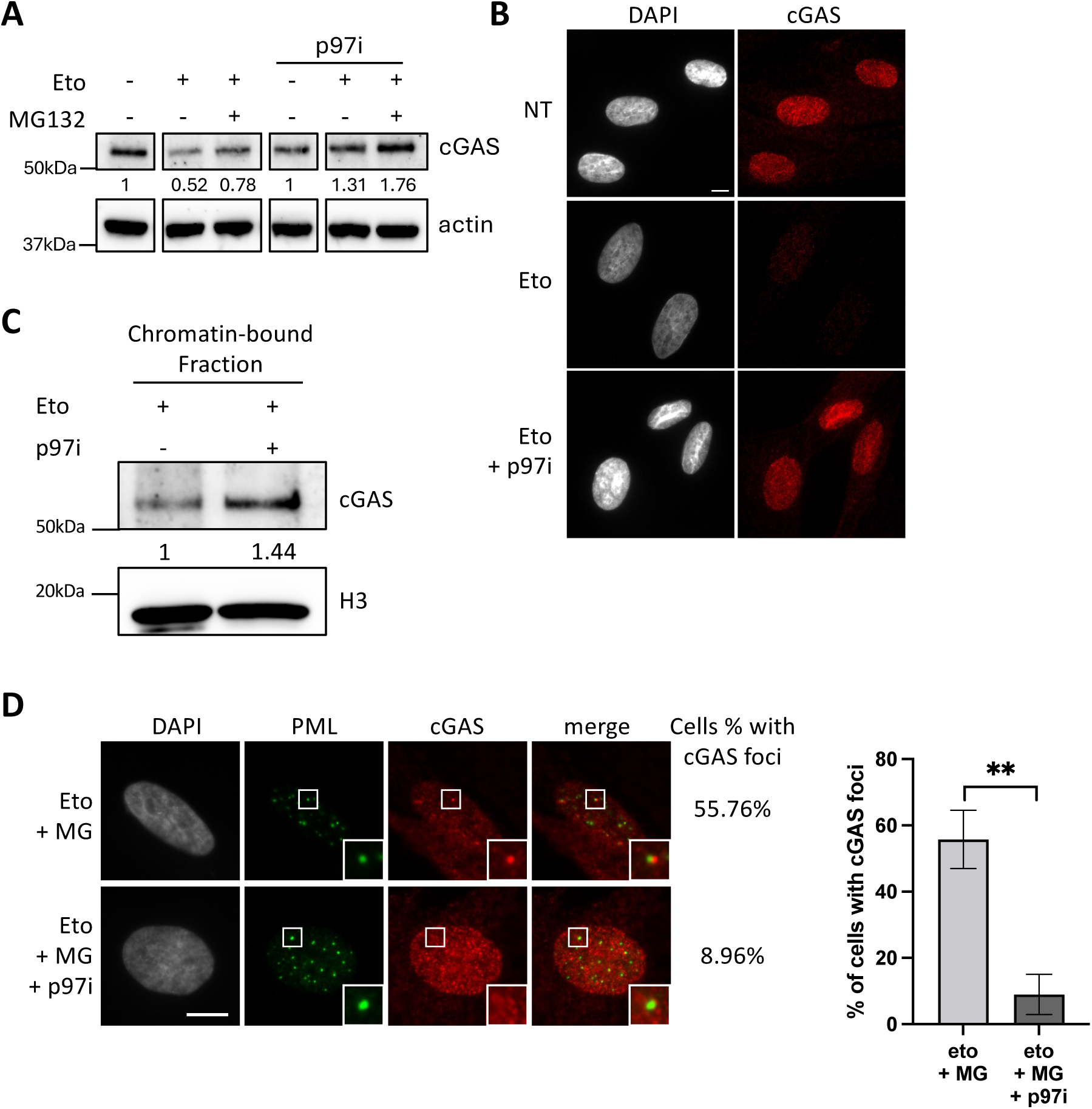
p97 is required for cGAS untethering from chromatin and allows cGAS foci formation. (A) Western-blot visualization of cGAS from total cell extracts on BJ cells treated with etoposide at 10μM for 24h, p97i (CB5083) at 1μM for 24h and MG132 at 2,5μM for 8h. Actin is a loading control and quantification of cGAS levels relative to actin are shown below the WB (numbers are representative from 2-3 independent experiments). (B) Fluorescence microscopy visualization of cGAS (red) in BJ cells treated with etoposide at 10μM for 24h +/- p97i (CB5083) at 1μM for 24h. Scale bar is 10μm. (C) Western-blot visualization of cGAS from the chromatin-bound fraction of BJ cells treated as in (B). H3 is a loading control and quantification of cGAS levels relative to H3 are shown below. (D) (Left) Fluorescence microscopy visualization of PML (green) and cGAS (red) in BJ cells treated as in (A). Cell nuclei are visualized by DAPI staining (grey). Scale bar is 10μm. Insets represent enlarged images (3X) of selected areas. (Right) Histogram shows quantitative analysis of cells with cGAS foci juxtaposing PML NBs (n=3 independent experiments). P-values (Student t-test) : **<0,01.

### cGAS preferentially accumulates in heterochromatin regions and spreads onto euchromatin regions upon proteasome inhibition

Based on the observations that cGAS is released from chromatin upon etoposide treatment but remains partially chromatin-bound upon proteasome inhibition, we next investigated the genome-wide distribution of endogenous cGAS by CUT&RUN under these conditions. As a negative control, we included a non-targeting IgG, and as a positive control, H3K4me3. As expected, peak annotation revealed that H3K4me3 was almost exclusively enriched on gene promoters (Figure 6A and Sup. Figure 6A), with a characteristic dip in signal at the nucleosome-free region immediately upstream of the TSS, while no enrichment was detected with the negative control. In contrast, cGAS localized predominantly to intronic and distal intergenic regions in untreated cells (Figure 6A) and showed no accumulation on H3K4me3-rich peaks (Sup. Figure 6B). Given that exogenous GFP-cGAS has been reported to associate with H3K9me3-rich heterochromatin regions (including LINE elements), and α-satellite centromeric regions ^33^, we next examined whether endogenous cGAS displays similar localization. Using published H3K9me3-enriched regions ^40^, we observed robust cGAS enrichment on these heterochromatin regions (Figure 6B), as well as at KRAB domain-containing zinc finger proteins (KZFP) encoding genes (Figure 6C), which are organized in highly repetitive gene clusters ^41^. For instance, cGAS was strongly enriched on two KZFP clusters on chromosome 19 (Figure 6D and Sup. Figure 6C), whereas no enrichment was observed at neighboring non-KZFP genes on the same chromosome (Sup. Figure 6D). Because etoposide treatment reduces the pool of chromatin-bound cGAS (Figures 1C-D), we asked whether this was reflected in a loss of cGAS in its enriched regions. However, no significant decrease was detected, suggesting that CUT&RUN predominantly captures few high-affinity cGAS binding sites, that remain unaffected by etoposide treatment (Figures 6B-E). In contrast, MG132 treatment in the context of DNA damage markedly altered cGAS distribution: binding paradoxically decreased slightly on heterochromatin regions (Figure 6B) and KZFP genes (Figures 6C-D), while spreading beyond KZFP loci (Sup. Figure 6E) suggesting a redistribution of the chromatin-bound pool of cGAS. Genome-wide peak annotation further confirmed this redistribution toward euchromatin, with increased cGAS occupancy at gene promoters (Figure 6A). Notably, cGAS became significantly enriched on LINE elements, which are interspersed in euchromatin regions (Figures 6F-H), as well as on ERV elements (Sup. Figure 6F). Together, the CUT&RUN data highlight that, while cGAS preferentially associates with specific heterochromatin sites, it expands into euchromatin regions, if not degraded following DNA damage and proteasome inhibition.

**Figure 6:**
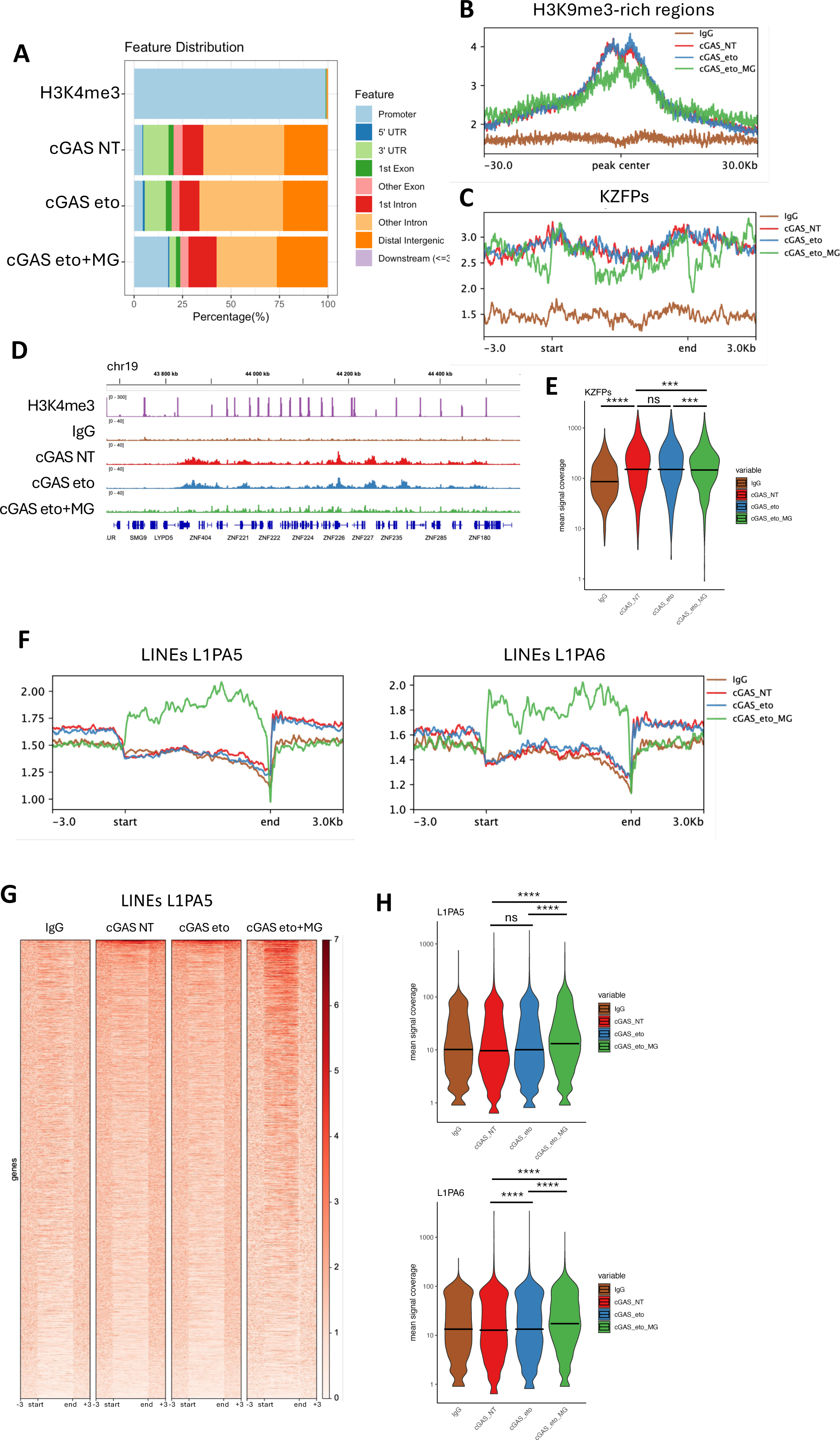
cGAS is enriched on heterochromatin but spreads on euchromatin regions in conditions of DNA damage upon proteasome inhibition. (A) CUT&RUN against endogenous cGAS or H3K4me3 was performed in BJ primary cells treated with etoposide at 10μM for 24h +/- MG132 at 2,5μM for the last 8h. Plot shows the genomic feature distribution of identified peaks using MACS2 in the different conditions. (B) Profile plots of cGAS density on H3K9me3-rich regions identified in ^40^. (C) Profile plots of cGAS density on KZFP genes listed in ^41^. (D) Genome browser snapshot of IgG, H3K4me3 and cGAS enrichment in BJ cells treated as in (A), across the indicated region. (E) CUT&RUN mean signal coverage of cGAS on KZFPs across the different samples. Adjusted p-values: *** <0.001; **** <0.0001 (Wilcoxon rank-sum test), ns: non-significant. (F) Profile plots of cGAS density on specific families of LINE-1 (L1PA5 and L1PA6). (G) Heatmaps showing the density of cGAS signal on L1PA5 LINE-2 subfamily. (H) CUT&RUN mean signal coverage of cGAS on L1PA5 or L1PA6 across the different samples. Adjusted p-values: **** <0.0001 (Wilcoxon rank-sum test), ns: non-significant.

### Accumulation of cGAS on chromatin correlates with an impaired DNA damage response and senescence entry

Given the importance of the DNA damage response for senescence induction ^42^, and building on our observations of cGAS nuclear dynamics, we next investigated their functional relevance in this process. Specifically, we first asked whether the nucleoplasmic pool of cGAS, independently of STING, contributes to early senescence entry, and second, whether blocking its release from chromatin affects the DDR or senescence induction. To address these questions, we first investigated which of the hallmarks of senescence ^43^ could be detected in BJ cells, as early as after 24h of etoposide treatment. As expected, a global loss of proliferation was observed via EdU-incorporation at day 1 post-etoposide treatment, which remained at day 5 or 7 post-etoposide (Sup. Figure 7A), together with SASP induction (Sup. Figure 7B). Cells were also positive for the senescence associated beta-galactosidase activity (SA-β-gal) at day 10 post-treatment (Sup. Figure 7C). The accumulation of the histone chaperone HIRA in PML NBs, which is another known marker of senescence required for senescence associated heterochromatin foci (SAHF) formation ^44,45^ and SASP induction ^46^, could be detected as early as 24 hours post-treatment in about 48% of the cells, prior to the onset of the other senescence markers and concomitant with γH2A.X induction (Figure 7A). Its frequency further increased during a time-course recovery (Sup. Figure 7D), while DNA damage itself did not trigger HIRA accumulation in PML NBs (Sup. Figure 7E and 47). RNA-seq analysis of BJ fibroblasts treated with etoposide for 24h revealed widespread, time-dependent transcriptional changes persisting at 72h of treatment (Sup. Figure 7F). GO analysis showed downregulation of cell cycle and DNA replication genes, and upregulation of p53 signaling, apoptosis, and autophagy pathways, consistent with early senescence markers at 24h post-etoposide (Sup. Figure 7F.)

**Figure 7:**
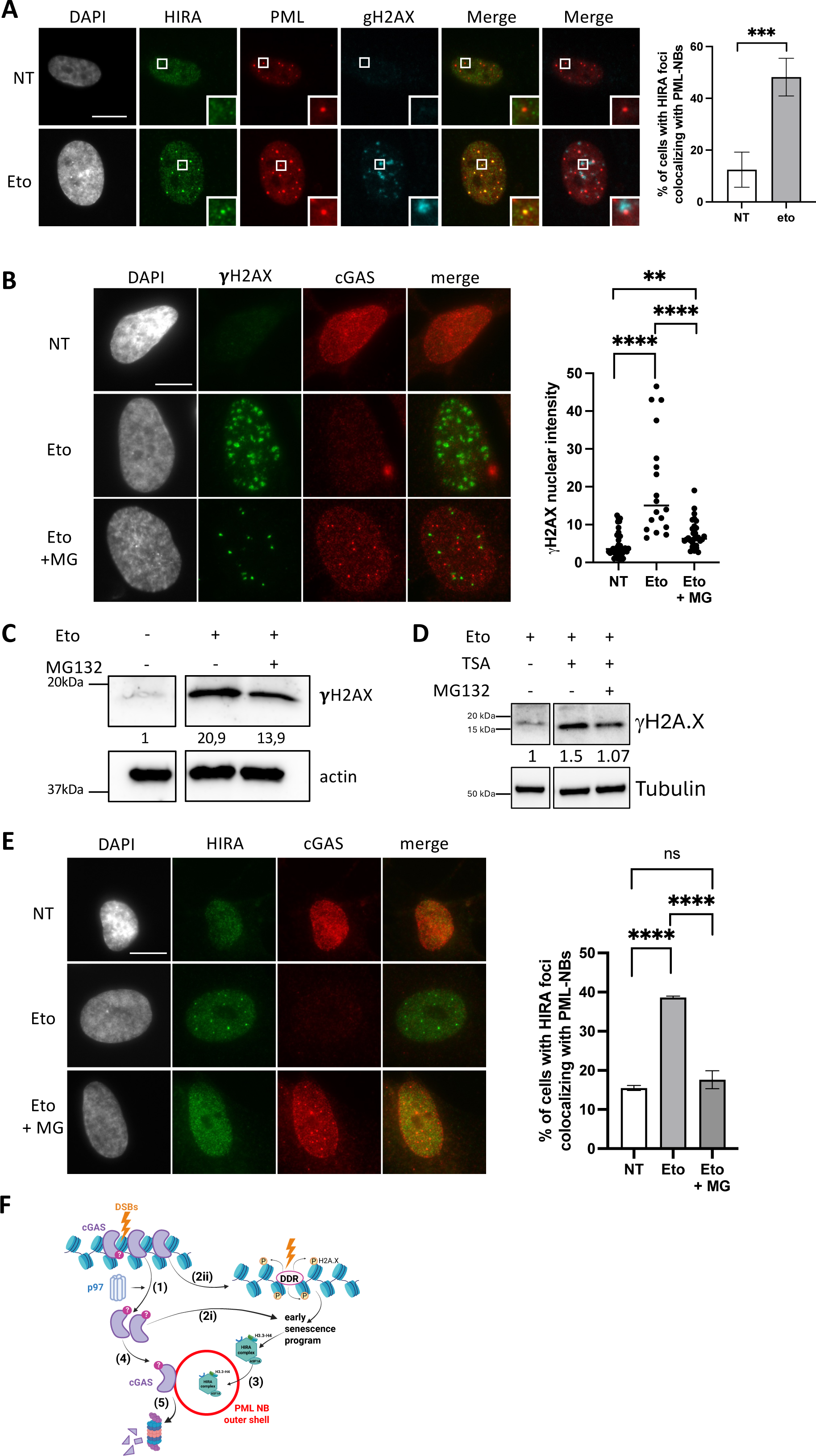
Inhibition of cGAS degradation correlates with a decreased DDR and impaired senescence entry. (A) (Left) Fluorescence microscopy visualization of HIRA (green), PML (red) and γH2AX (cyan) in BJ cells treated with etoposide at 10μM for 24h. Cell nuclei are visualized by DAPI staining (grey). Scale bar is 10μm. Insets represent enlarged images (3X) of selected areas. (Right) Histogram shows quantitative analysis of cells with HIRA localization at PML NBs (n=3 independent experiments). P-values (Student t-test) : ***<0,001. (B) Fluorescence microscopy visualization of γH2AX (green) and cGAS (red) in BJ cells treated etoposide at 10μM for 24h and with MG132 at 2,5μM for 8h. Cell nuclei are visualized by DAPI staining (grey). Scale bar is 10μm. (Right) Plot shows quantitative analysis of nuclear γH2AX intensity in cells. P-values (Student t-test) : **<0,01 ; ****<0,0001. (C) Western-blot visualization of γH2AX from total cell extracts on BJ cells treated with the same conditions as in (A). Actin is a loading control and quantification of γH2AX levels relative to actin are shown below the WB (numbers are representative from three independent experiments). (D) Western-blot visualization of γH2AX from total cell extracts on BJ cells treated with etoposide at 10μM for 24h, TSA at 2μM for 24h and MG132 at 2,5μM for 8h. Tubulin is a loading control and quantification of γH2AX levels relative to tubulin are shown below the WB (numbers are representative from one experiment). (E) (Left) Fluorescence microscopy visualization of HIRA (green) and cGAS (red) in BJ cells treated with etoposide at 10μM for 24h and with MG132 at 2,5μM for 8h. Cell nuclei are visualized by DAPI staining (grey). Scale bar is 10μm. (Right) Histogram shows quantitative analysis of cells with HIRA localization at PML NBs (n=3 independent experiments). P-values (Student t-test) : ****<0,0001, ns: non significant. (F) Putative model for nuclear cGAS dynamics upon DNA damage in primary cells. Upon DSB induction, cGAS might be modified and is untethered by p97 (1). Free nucleoplasmic cGAS participates in the early senescence entry independently of STING (2i). In addition, cGAS untethering from chromatin correlates with establishment of a complete DDR (DNA damage response) (2ii) which participates in the early senescence program, as exemplified by HIRA accumulation in PML NBs (3). The extra-pool of nucleoplasmic cGAS localizes next to PML NBs (4) which participates in its degradation in a proteasome- and Cullin3-dependent manner (5), thus balancing cGAS nucleoplasmic levels with its function in early senescence entry.

We therefore asked whether nuclear cGAS is required for early senescence entry, independently of STING. While the cGAS-STING pathway is key to promote the senescence-associated secretory phenotype (SASP) in late senescence ^6,7,48^, the STING-independent contributions of nuclear cGAS to the early stages of senescence remain unclear ^48^. Using siRNAs to deplete cGAS or STING (Sup. Figure 7G) prior to 24h of etoposide treatment, we monitored HIRA accumulation in PML NBs. Loss of cGAS severely impacted this early senescence marker, whereas STING depletion had no effect (Sup. Figures H-I). These data agree with a recent report also describing a STING-independent role of nuclear cGAS which, itself, contributes to activation of SASP genes ^49^.

To assess the potential significance of cGAS release from chromatin, we examined whether blocking its degradation by proteasome inhibition, resulting in a global increase of cGAS on chromatin (Figure 2B) and its redistribution to euchromatin regions (Figure 6A), would impact the DNA damage response (DDR) and the onset of early senescence. Cells treated with etoposide together with MG132, showed diminished γH2A.X levels, as shown by immunofluorescence and western blot (Figures 7B-C). The lower amount of γH2A.X foci and their smaller size was intimately linked with higher cGAS nuclear levels and cGAS foci formation (Figure 7B). Treatment with p97i concomitant with etoposide had similar effects, blunting the amount of γH2A.X, which anti-correlated with cGAS amounts in the nucleus (Sup. Figure 8A) and on chromatin (Figure 5C). p97i effect was more drastic than the effect of MG132 on dampening γH2A.X levels, possibly owing to the fact that, for toxicity reasons, MG132 was only added in the last 8 hours of etoposide treatment, when a pool of cGAS is already being detached from chromatin, allowing the formation of small γH2A.X foci. Notably, no cGAS foci were detected after p97i, consistent with our earlier observations that p97i prevents cGAS unhooking from chromatin (Figure 5D), which may therefore hinder cGAS foci formation. In contrast, treatment with TSA, which triggers cGAS removal from chromatin, led to a significant increase in γH2A.X levels (Figure 7D). Collectively, these results suggest that stabilizing cGAS on chromatin or preventing its degradation correlates with a dampened DNA damage response, whereas its removal from chromatin enhances it. We then investigated whether preventing cGAS degradation blunts early senescence entry by quantifying the amount of cells with HIRA accumulation in PML NBs. Addition of MG132, which increased the pool of nuclear cGAS, prevented HIRA relocalization in PML NBs, suggesting an impaired senescence entry (Figure 7E). Together, these findings demonstrate that the DNA damage-induced chromatin destabilization triggering cGAS unhooking from chromatin correlates with early senescence induction. The p97-PML NBs axis might ensure the control of nucleoplasmic cGAS abundance to limit a potential cell-intrinsic activation of innate immune responses.

## Discussion

By assessing the dynamics of nuclear cGAS following DNA damage, we provide important novel insights into the mechanisms controlling cGAS levels and role in the nucleus. We explore how these mechanisms might influence therapeutic strategies targeting cGAS.

### Chromatin-bound cGAS is released upon chromatin destabilization and degraded through a p97-PML NBs axis

While it is now well-established that cGAS is a nuclear protein, tightly bound to chromatin to remain inactivated ^10,11^, our data show that, upon DNA damage, a fraction of cGAS is unhooked from chromatin by p97 followed by its degradation through a PML NBs-Cullin3-proteaxome axis. In this context, post-translational modifications (PTMs) of proteins, such as SUMOylation and ubiquitination, have emerged as significant mechanisms for regulating diverse aspects of the DDR, orchestrating the recruitment/turnover of DDR proteins at DNA damage sites which is essential for preserving genome integrity ^50^. While our attempts have remained so far unsuccessful in the identification of cGAS SUMOylation, probably owing to the low levels of cGAS in BJ cells, several screens have identified the lysine K285 as a target for SUMOylation ^51,52^. Interestingly, both the SUMO E2 conjugating enzyme UBC9 and the SUMO E3 ligases PIAS1 and PIAS4 accumulate at sites of DSBs where they are required for accrual of DDR factors and proficient DSB repair ^53,54^. In addition, proteins belonging to the class of STUbLs (SUMO-Targeted Ubiquitin Ligases), such as RNF4 and RNF111 which can ubiquitinate SUMOylated proteins connecting them to the proteasome-dependent degradation pathway, are also recruited to DSBs where they regulate protein turnover ^55–58^. Remarkably, all these actors of the SUMOylation/ubiquitination pathways localize both to DSBs as well as in PML NBs ^28,59,60^ underscoring the tight interplay between PML NBs and the turnover of proteins during the DDR. Our data indeed suggest the existence of two different pools of PML NBs, one implicated in the DNA damage response and another one dealing with the undegraded nucleoplasmic cGAS (Sup. Figure 2E). PML NBs, by simultaneously concentrating enzymes and their substrates within a specific oxidation-protective environment, distinct from the rest of the nucleoplasm, could thus orchestrate the SUMOylation/ubiquitination of specific proteins regulating their stability ^27,61^. It is plausible that cGAS undergoes modification directly at DSBs, is subsequently extracted from chromatin by p97 (see below), and then targeted to PML NBs for degradation. Of note, using JASSA ^62^, we found 2 predictive SIM sites in cGAS (aa 151-154 and aa 308-311), which may play a role in cGAS recruitment adjacent to PML NBs. Alternatively, p97 could directly bring cGAS to PML NBs since it localizes in these bodies in a SUMOylation dependent manner ^28^. Altogether, SUMOylation remains a key regulator of cGAS nuclear dynamics upon DNA damage, involved in cGAS targeting to PML NBs followed by its degradation (Figures 3E-H). Whether other cGAS modifications, such as phosphorylation on Serine 37 that regulates cGAS chromatin tethering ^63^, are implicated in this context remains to be determined.

Proteasome inhibition after DNA damage results in two distinct cGAS pools: an increased chromatin-bound fraction, which may reflect either cGAS that remained tethered or released cGAS that rebound to chromatin, and a nucleoplasmic fraction that accumulates in foci adjacent to PML NBs. cGAS foci are reminiscent of the ones formed by KAP1, an essential repressor of the transcription of transposable elements, which is SUMOylated by PML and accumulates in foci adjacent to PML NBs upon proteasome inhibition ^61^. Remarkably, KAP1 which is phosphorylated upon DNA damage and plays essential role in DSB repair, is targeted for degradation by RNF4, in cooperation with the p97, to allow for HR repair ^64,65^. Our data show the essential role of p97 for cGAS unhooking from chromatin and its accumulation in foci adjacent to PML NBs upon proteasome-inhibition. We cannot exclude that cGAS present in foci adjacent to PML NBs is chromatin-bound, but it seems unlikely given that (i) cGAS would accumulate on chromatin at only a dozen of particular sites which is inconsistent with our CUT&RUN data, and (ii) the requirement for p97 to form cGAS foci suggests that it represents a nucleoplasmic pool of undegraded cGAS.

Upon dsDNA binding, multivalent interactions of cGAS with DNA and other cGAS molecules drive the formation of phase-separated liquid droplets that concentrate cGAS with DNA in a crowded cytoplasm, which is critical to restrict DNA degradation and allows efficient innate immune activation ^66–68^. Formation of cGAS foci in the nucleoplasm upon proteasome inhibition may resemble the formation of liquid-like droplets thanks to multivalent interactions of cGAS. Yet, we hypothesize that these foci of undegraded nucleoplasmic cGAS are kept in check by PML NBs to avoid its overt activation. While PML NBs have intimately been linked with proteasomal degradation ^69^, we confirm here a close connection between an important 19S component, ADRM1, PML NBs and cGAS in conditions of DNA damage and proteasome inhibition (Figure 3A). Interestingly, proteasome foci containing ADRM1 and active 26S proteasomes, recently characterized in senescent cells as SANPs (Senescence Associated Nuclear Proteasome foci), were shown to colocalize with p97 ^37^. We further reveal the importance of Cullin3 in targeting cGAS for degradation through PML NBs. While removal of the SPSB3-CRL5 degradation complex of cGAS ^22^ had no impact on cGAS nuclear dynamics, this suggests the existence of complementary pathways specific for the degradation of nuclear cGAS following acute stresses. Altogether, our data extend the importance of PML NBs for controlling nuclear levels of proteins involved in essential DNA transactions ^70,71^.

### Chromatin-bound cGAS untethering correlates with the DNA damage response activation and senescence entry

Our data show that cGAS is released from chromatin and degraded after DNA damage, which may seem contradictory with earlier studies showing cGAS accumulation at DNA damage sites. Indeed, cGAS is recruited to DNA damage sites at early times after damage and abrogates the formation of a functional PARP1-Timeless complex required for homologous recombination (HR) ^19^. In addition, cGAS also negatively regulates HR through DNA compaction which inhibits the formation of the Rad51 filament for homology searching ^18^. We hypothesize that cGAS could be recruited transiently at DNA breaks, followed by its rapid degradation to allow the DNA damage response ^18^. This hypothesis is consistent with the fact that we observe release of only a fraction of the chromatin-bound cGAS pool, which could mainly consist of cGAS present at DNA damage sites, coupled with a pool of cGAS resulting from its cell-cycle dependent turnover from chromatin ^22^.

Chromatin opening is required for γH2A.X spreading as shown with the decondensation of heterochromatin regions upon DSBs ^72^. Here, by using MG132 which prevent degradation in conditions of DNA damage, we show an impaired formation of γH2AX, correlating with an increase of cGAS on chromatin, leading to an altered DNA damage-induced senescence entry. In addition, inhibition of p97, which prevents cGAS release from chromatin, also blunted formation of γH2A.X foci (Sup. Figure 8A). Interestingly, depletion of p97 leads to reduced protein turnover at DNA damage sites, and abolishes BRCA1 or Rad51 recruitment ^73^, which is inhibited by cGAS ^18^. Conversely, by using TSA, which drastically releases cGAS from chromatin, we correlate the absence of cGAS from chromatin with a strong γH2A.X signal seen upon combined TSA+etoposide treatment as compared to etoposide alone. Thus, our data are consistent with a role of cGAS as a negative regulator of the DDR adding the requirement of cGAS untethering from chromatin to allow an efficient DDR and senescence entry. It is also interesting to note that cGAS is mainly enriched at the 3ʹ end of ZNFs (Sup. Figure 6C). This particular region of ZNFs is H3K9me3-rich and has been suggested to create a more condensed chromatin structure to inhibit homologous recombination between highly similar ZNF genes ^74^. cGAS binding on 3’end of ZNFs could thus also contribute to inhibit recombination of these repeated regions. Our CUT&RUN data showing cGAS spreading onto euchromatin and gene promoters is also intriguing in regards to a recent report uncovering cGAS-dependent recruitment of the SWI/SNF chromatin remodeler complex to specific regions ^63^. While the role of cGAS in transcriptional regulation remains unexplored, it is tempting to speculate that the broad spreading of cGAS on chromatin observed in conditions of DNA damage and proteasome inhibition, could also alter the transcriptional response required for senescence entry.

### Fine-tuning nucleoplasmic/chromatin-bound cGAS levels is essential to regulate senescence entry

Previous studies have underscored a role of cGAS-STING in the late induction of the SASP during senescence ^6,7,48^. In addition, recent data uncover a STING-independent function of cGAS which contributes to SASP induction in a DNA demethylation model of senescence induction ^49^. Here, we confirm a key role of cGAS, independent of STING, in promoting early senescence entry in a DNA damage-induced senescence model (Sup. Figures 7G-I). Yet, we also show that degradation of cGAS, after induction of DNA damage, is required for early senescence induction upon DNA damage induction, which might seem paradoxical. A few hypotheses can help to resolve this paradox: first, DNA damage only induces a partial degradation of the chromatin-bound/nucleoplasmic pool of cGAS, leaving a pool of nucleoplasmic cGAS available to perform various functions independently of STING. Interestingly, following herpes simplex virus 1 infection, cGAS is released from chromatin in the nuclear soluble fraction, which activates cGAMP production following viral DNA sensing ^75^. Second, the dynamics of cGAS release from chromatin itself could participate in the DDR response and help trigger a senescence-inducing response. Third, we cannot exclude the existence of, as yet to be discovered, specific nuclear functions for cGAS in senescence, which may relate to transcriptional regulation ^63^.

We thus propose the following model in which the fine-tuning of cGAS levels through a p97-PML NBs axis is key to allow senescence entry and, at the same time, avoid an overt activation of cGAS: following DNA damage, a portion of cGAS is untethered from chromatin allowing an efficient DNA damage response and triggering early senescence entry. Part of the detached pool of nucleoplasmic cGAS could also participate in launching an early senescence program independently of STING (Sup. Figures 7G-I), possibly by contributing to SASP expression ^49^, yet the extra pool of cGAS is subsequently degraded to avoid unintended innate immune activation. cGAS might be SUMOylated and/or ubiquitinated directly on chromatin, which is the signal for p97 to segregate it from chromatin. cGAS SUMOylation/SIM would ensure its targeting next to PML NBs, that ensure its degradation through their intimate connection with the proteasome-degradation pathway (Figure 7F). Preventing cGAS dynamic untethering from chromatin correlates with a dampened DNA damage response and impaired early senescence entry.

Activation of the cGAS-STING pathway has contrasting effects on cancers occurrence and progression. While cGAS-STING signaling is beneficial for antitumor immunity, the inflammatory cytokine secretion may fuel cancer progression or metastasis, suggesting that the effects of the cGAS-STING pathway could be context- and stage-dependent in various cancers. Our data underline an important difference between the response of primary cells and cancerous cells in the regulation of cGAS nuclear levels upon DNA damage, which could explain why cancerous cells fail to enter into senescence upon a DNA damage inducer such as etoposide. Together, our data underscore the critical importance of fine-tuning cGAS nuclear dynamics, whose specific STING-independent roles warrant further investigation.

## Methods

### Cell lines and lentiviruses production

Human BJ primary foreskin fibroblasts (ATCC, CRL-2522) and HeLa cells, were cultivated in DMEM medium (Fisher scientific, 41966) containing 5% of fetal bovine serum (FBS) (Fischer scientific, 10453302), 1% of penicillin/streptomycin (Sigma-Aldrich, P4458), at 37 °C under 5% CO2 and humid atmosphere. All cell lines were tested negative for mycoplasma contamination. Drugs and molecules used for cell treatments are described in the Key Resources Table in Appendix (duration is mentioned in the main text).

### siRNAs

BJ cells were transfected with 40nM of human siRNA for different timings (indicated in the main text for each experiment) using Lipofectamine RNAiMax reagent (Invitrogen, 13778–075) and Opti-MEM medium (Gibco, 31-985-070). siRNAs used and their sequences are summarized in the Key Resources Table in Appendix.

### Antibodies

All the primary antibodies used in this study, together with the species, the references and the dilutions for immunofluorescence and western blotting, are summarized in the Key Resources Table in Appendix.

### Immunofluorescence (IF)

Immunofluorescence was performed as in ^76^. Briefly, cells were seeded on coverslips in 6- or 24-well plates. After treatment, cells were washed with PBS (EuroBioScience, CS0PBS01) and fixed in paraformaldehyde 2% (Euromedex, EM-15735). Cells were permeabilized with PBS 0,2% Triton X-100 (Sigma-Aldrich, T8787) for 5min and washed in PBS 2 times before saturation in a 5% BSA solution (Dutcher, P06-1391100) diluted in PBS 0,1% of Tween (Sigma-Aldrich, P9416), written PBST. Cells were incubated in primary antibodies diluted in PBST with 5% of BSA for 1h (see the Key Resources Table in Appendix for antibodies dilution). Highly cross-absorbed goat anti-mouse or anti-rabbit (H+L) Alexa-488, Alexa-488 IgG1, Alexa-555, Alexa-594 IgG2b or Alexa-647 (Invitrogen Life Technologies) were used as secondary antibodies, which were incubated 30min in PBST 5% BSA. For triple staining immunofluorescence of cGAS-PML-ADRM1, cells were incubated with the cGAS and PML primary antibodies followed by washes, and incubation with the secondary antibodies as indicated above. Cells were then blocked again in PBST 5% BSA for 20min at R°C. ADRM1 antibody, directly coupled to CoraLite-Plus 555 fluorochrome using the FlexAble 2.0 kit (Proteintech #KFA502), was then incubated overnight at 4°C. Cells were then incubated in DAPI (Invitrogen Life Technologies, D1306) diluted in PBS at 0.1 μg/mL for 5min at RT°C. Coverslips were mounted in Fluoromount-G (SouthernBiotech, 0100–01) and stored at 4°C before observation.

### EdU incorporation and detection

Proliferating and senescent BJ cells were seeded on coverslips in 6-well plates. To label proliferating cells, cells were incubated with EdU (ethynyl deoxyuridine) at 20uM, an analog of thymidine, for 2h at 37°C. EdU detection was performed using the Click-iT EdU Alexa Fluor 555 imaging kit (Invitrogen, C10338). Cells were imaged as mentioned in microscopy part.

### Senescence assay

For senescence detection, cells grown on coverslips, were fixed with 3.7% formaldehyde for 3min and washed 3 times in PBS. SA-βGal activity was detected by incubating cells overnight at 37°C (without CO^2^) in a staining solution containing 1mg/mL of X-Gal (Sigma, B4252), 40mM citric acid/sodium phosphate buffer at pH=6, 5mM potassium ferricyanide, 5mM potassium ferrocyanide, 150mM NaCl and 2mM MgCl2. The cells were then washed 3 times with PBS and mounted in Fluoromount-G (SouthernBiotech, 0100–01) and stored at 4 °C before observation. Images were acquired on the Axioskop (Zeiss) upright and a color camera AxioCam ICc5 (Zeiss).

### Microscopy, imaging, and quantification

Images were either acquired with an Axio Observer Z1 inverted wide-field epifluorescence microscope (100 X or 63 X objectives/N.A. 1.46 or 1.4) (Zeiss) and a CoolSnap HQ2 camera (Photometrics) or an ORCA-Flash4.OLT3 CMOS camera (Hamamatsu). Identical settings and contrast were applied for all images of the same experiment to allow data comparison. Raw images were treated with Fiji software. HIRA complex accumulation in PML NBs and cGAS foci were attested by manual counting of a minimum of 100 cells for each condition and per replicate. PML-NBs and cGAS foci proximity was measured using the Fiji RenyiEntropy mask on PML and cGAS staining. X and Y coordinates for the center of the spots were recovered and all distances between each PML NB and cGAS foci were calculated using the formula d =√((x1-x2)^2^ + (y1-y2)^2^) to find the minimal distance in each nucleus. Quantification of nuclear intensities was performed with Fiji. To quantify cGAS intensity in the nucleus, we created a mask of nuclei based on DAPI staining threshold (Image calculator function of Fiji) and then applied the measure function on cGAS channel.

### Immunofluorescence for dSTORM imaging and dSTORM acquisition

Cells were seeded on Willco GWST 35–22 for dSTORM. 1h before fixation, cells were treated with SPY555-DNA (SC201 SPIROCHROME). Cells were fixed for 10min with 2% paraformaldehyde, quenched with 1M Tris for 10min, then permeabilized twice 5min in a PBS solution containing 0.5 % Triton X-100. Slides were saturated with 3% Normal goat serum (NGS) in PBS for 1h at RT. Primary antibodies were diluted in PBS containing 3% NGS. After incubation at 37°C for 2h, coverslips were washed three times with PBS containing 0.5% Triton X-100, and then incubated with cross-adsorbed goat anti-mouse F(ab’)2-FITC antibody (Bethyl, A90-241F) and lama-anti-rabbit(H + L)-CF-647 antibody (Ozyme, 20453). After a further 45min incubation at 37°C, Willco dishes were mounted with Eternity buffer (Everspark 2.0, Idylle) ^77^ and air-tight sealed with twinsil for dSTORM imaging (see details in section below). Samples were stored in the dark and at 4°C before observation.

Image acquisition or/and image analysis were performed on INMG-PGNM (CNRS UMR5261 INSERM U1315, Claude Bernard University, VXL super-resolution Service) with a Bruker Vutara VXL microscope. 3D super-resolution images were acquired based on single molecule localization (SML) biplane technology (Mlodzianoski et al., 2009), using a ×60-magnification,

1.3 NA silicon immersion objective (Olympus UPLSAPO60XS2) and an sCMOS ORCA BT fusion camera (Hamamatsu). A collection of 10’000 images by (50 µm^2^) was sequentially recorded after pumping steps for cGAS, SPY555-DNA and PML labelling using 638nm followed by 555nm and 488 excitation lasers at 10% (i.e. 1,856 kW/cm^2^), 60% (i.e. 10,56 kW/cm^2^) and 10% (i.e. 4,08 kW/cm^2^) respectively with an exposure time of 10msec, 20msec and 20msec. Gaussian fitting with a background level of 20 for cGAS and SPY555-DNA and 10 for PML was directly performed with the Vutara SRX software to identify the x,y fluorescence profile of each blink on both planes. Adequation with a reference collection of profile pairs from the 3D experimental point-spread function (PSF) measured from calibrated biplane data using 100nm diameter Tetraspeck beads (Thermo Fisher Scientific T7279), allowed to assess z coordinate for each blink and perform 3D reconstruction over a 900nm thickness volume. Cluster analysis was then calculated with the DBSCAN algorithm implemented in SRX software, using a maximum particle distance of 100nm, a minimum particle count to form cluster of 10 and a hull alpha-shape radius of 100nm.

### Western blotting (WB)

Total cellular extracts were obtained by directly lysing the cells in 2 X Laemmli sample buffer (LSB) (125 mM Tris-HCl pH 6.8, 20% glycerol, 4% SDS, bromophenol blue) containing 100 mM DTT. Samples were boiled for 10min and loaded on 7,5 or 10% SDS-polyacrylamide gels (Biorad, 161-0181 and 161-0183). After electrophoresis, proteins were transferred on nitrocellulose membranes using a Transfer kit (Biorad, 1704271) in the Biorad Trans-Blot Turbo apparatus. Membranes were saturated in PBST with 5% of milk for 30 min and then incubated in primary antibody diluted in PBST 5% BSA for 1 h at room temperature or at 4°C overnight. After 3 washes in PBST, membranes were incubated with secondary antibodies diluted in PBST (see the Key Resources Table in Appendix for antibodies dilution). Signal was revealed on ChemiDoc Imaging System (Bio-Rad) by using Amersham ECL Prime Western Blotting Detection Reagent (GE Healthcare Life Sciences, RPN2236) or Clarity Max Western ECL Blotting Substrate (Bio-Rad, 1705062).

#### Fractionation

Subcellular fractionation of proteins was performed using the Subcellular Protein Fractionation Kit for Cultured Cells (Thermo Fisher Scientific, #78840). Briefly, the cells were washed once with ice-cold PBS. After completely removing PBS, the cells were collected in cold PBS with 1 mL per well in a 6 wells plate. After PBS removal the cells were dissolved using 200uL of CEB solution. The lysates tubes were incubated for 10min at 4°C. The supernatants were collected as cytoplasmic fraction after centrifugation at 500g for 5min. The cell pellets were dissolved using 200 μL of MEB solution. After a 5 s vortex, the lysate tubes were incubated for 10 min at 4°C. The supernatants were collected as membrane fraction after centrifugation at 3000 x g for 5 min. The pellets were dissolved using 100uL of NEB solution. After 15s vortex, the lysates tubes were incubated for 30 min at 4°C. The supernatants were collected as nuclear soluble fraction after centrifugation at 5000g for 5 min. The pellets were dissolved using 100uL of NEB complemented solution. After 15s vortex, the lysates tubes were incubated for 15 min at room temperature. The supernatants were collected as chromatin-bound fraction after centrifugation at 16000g for 5min. Samples concentrations were determined by Bradford assay. Same amounts of lysates were diluted in 2X LSB containing 100mM DTT and loaded on gels.

### Reverse transcription (RT)

TRIzol reagent protocol (Invitrogen, 15596026) or RNeasy Mini kit (Qiagen, 74104) were used to isolate total RNAs, resuspended in ddH2O according to the manufacturer instructions. Contaminant DNA was removed with the RNAse-free DNase Ambion (Invitrogen, AM2222). We used 1 μg of RNA for reverse transcription (RT). RNAs were annealed with Random Primers (Promega, C118A) and RT was performed with the RevertAid H Minus Reverse Transcriptase (Thermo Scientific, EP0452) according to the manufacturer instructions. cDNAs were stored at –20 °C before qPCR analysis.

### Quantitative PCR (qPCR)

qPCRs were performed using the KAPA SYBR qPCR Master Mix (SYBR Green I dye chemistry) (KAPA BIOSYSTEMS, KK4618). Primers used for qPCR are described in the Key Resources Table in Appendix.

### RNA-seq

BJ cells were treated for 24h or 72h with etoposide at 10μM or left untreated. RNAs were collected with the RNeasy Mini kit (Qiagen, 74104) and sent to BGI for sequencing on a DNBSEQ. Standard bioinformatics analysis was performed by BGI with the version 38 of the human reference genome (hg38). Dr. Tom Data Visualisation Solution (BGI) was used to perform the KEGG enrichment analysis. Heatmaps were done in R.

### CUT&RUN

BJ cells were first treated for 24h with etoposide at 10μM +/- addition of MG132 at 2.5μM in the last 8h or left untreated. At the end of the treatment, 600 000 cells for each condition were detached by trypsin-EDTA incubation, washed with growth medium and pelleted at 600g for 3min. CUT&RUN was then performed using Active Motif’s ChIC/CUT&RUN Assay kit (#53180) following the manufacturer’s instructions. Nuclei were first isolated by resuspending pellets in nuclei isolation buffer and incubating them for 10 min on ice. Nuclei were then washed twice in Complete Dig-Wash buffer and bound to activated Concanavalin A beads for 10 min at room temperature. Following that, nuclei were incubated with antibodies overnight at 4°C : 1uL of the positive control Rabbit mAb Tri-Methyl-Histone H3 (Lys4) (C42D8 Cell Signaling #9751), 1uL of the negative control Rabbit mAb IgG XP® Isotype Control (DA1E Cell Signaling #66362 or 1uL of Rabbit mAb cGAS (D1D3G Cell Signaling #15102). After antibody incubation, cells bound to concanavalin beads were washed twice with Cell permeabilization buffer to remove unbound antibodies. 2.5uL of pAG-MNase was then added in a total volume of 50uL of Cell permeabilization buffer and samples were incubated for 1h at 4°C. Cells were washed twice with Cell permeabilization buffer then cooled to 0°C for 5min and incubated with ice-cold 100 mM CaCl2 for 2h at 4°C. MNase digestion was terminated with the addition of STOP buffer containing 0.2mg/mL of glycogen and 0.2mg/mL of RNAseA followed by an incubation for 30min at 37°C. Following RNA digestion, protein digestion was performed by incubating samples with 0.1% of SDS and 0.3 mg/ml of proteinase K for 2h at 65°C. DNA was then diluted in 100uL of TE and purified by two rounds of extraction with Phenol-chloroform-isoamylic alcohol (Sigma #77617) and chloroform. The extracted aqueous phase was supplemented with 200mM NaCl. DNA was precipitated by addition of 2 volumes of 100% EtOH and incubation for 1h at -80°C. DNA was pelleted at 16 000g for 20min at 4°C, washed with 1 volume of 70% EtOH and pelleted again. The DNA pellet was air-dried and dissolved in 30uL of molecular biology grade water and was stored at –20 °C before library preparation. The libraries were paired-end sequenced on a DNBSEQ Technology platform (BGI Genomics Co., Shenzhen, China).

### CUT&RUN analysis

After removing adaptor sequences and low-quality reads by cutadapt (version 4.6), CUT&RUN paired-end reads were aligned to the human genome (GRCh38) using Bowtie 2 (v2.5.4) with a --very-sensitive setting. Sequence alignment map files, including uniquely mapped and multi-mapped reads, were transformed and sorted into BAM files using SAMtools (v.1.9). Quality controls were done using deepTools (v3.5.4). Normalization factors were calculated using CUT&RUN greenlist following the protocol of ^78^. Briefly, read counts in the .bed greenlist regions were quantified using deepTools multiBamSummary (v3.5.4). Size factors were calculated using DESeq2 in Rstudio (v2025.05.0+496). These size factors were then used to convert the alignment files (BAM format) to read coverage files in Bigwig format using deepTools bamCoverage (v.3.5.4) with option --scaleFactor and a 10 bp window size. The normalized bigwig tracks were visualised with IGV (version 2.15.1). Peak calling was either done using SEACR v1.3 ^79^ with the following parameters: normalization (non), peak calling (stringent) and the normalized bedgraph files (control file : IgG and experimental file : cGAS or H3K4me3), or using MACS2 (v2.2.7.1) with the following parameters : -t experimental.bam -c IgG_control.bam -n sample.name --ratio (scale factor sample/scale factor control) -f BAMPE - g hs --keep-dup all -q 0.01. Peak annotation was performed in Rstudio (v2025.05.0+496) with the readPeakFile, annotatePeak and plotAnnoBar functions of the ChIPseeker package. Heatmaps and profile plots were generated with deepTools computeMatrix, plotHeatmap and plotProfile commands (v.3.5.4). We used a .bed file generated with Table S2 from ^40^, GSE63116) to plot cGAS signal on H3K9me3 rich regions that show decreased H3K9me3 levels as a result of loss of HUSH. KZFPs .bed file was generated from Table S1 from ^41^. The LINEs and ERVs sequences in a .bed format were downloaded from the RepeatMasker UCSC genome browser9 (GRCh38/hg38). Subtypes of LINEs or ERVs were filtered in Bash using the grep command to allow creation of heatmaps on specific subtypes of LINEs (eg L1PA6). The FeatureCounts command of the subread package (v2.0.6) was used to count reads on the LINEs or KZFPs .bed files converted into .saf files. After verifying that the data does not follow a normal distribution with a Kolmogorov-Smirnov test, we tested whether the mean signal coverage of two samples differ significantly from each other by using the non-parametric Wilcoxon rank-sum test. All bioinformatic analyses were performed on the Institut Français de Bioinformatique (IFB) core cluster through JupyterHub. Figures were made with RStudio (v2025.05.0+496) or with Inkscape (v.1.0.2 (e86c8708, 2021-01-15)).

### Statistical analyses and figures

Histograms and statistical analyses were performed using GraphPad Prism 10. To perform Student t test, we verified normal distribution of samples using Shapiro test and variance equality with Fisher test. Mann-Whitney u-test was applied in absence of normality for the sample distribution. p-Values are depicted on graphs as follows: *<0.05; **<0.01; ***<0.001; ****<0.0001. Biorender.com was used to generate figures schemes and model.

### Materials availability

All plasmids and cell lines generated in this study can be accessed upon request to the corresponding authors.

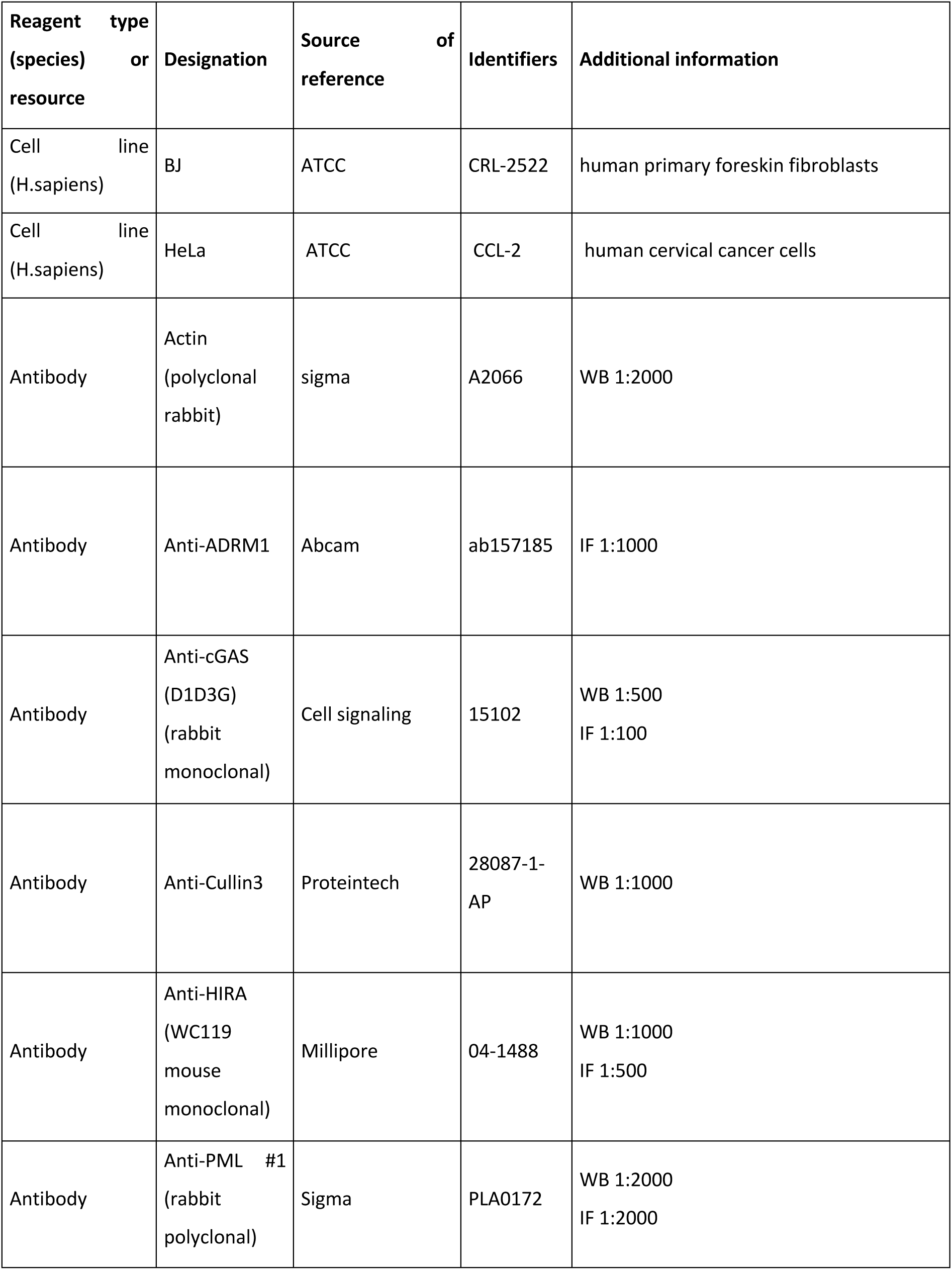
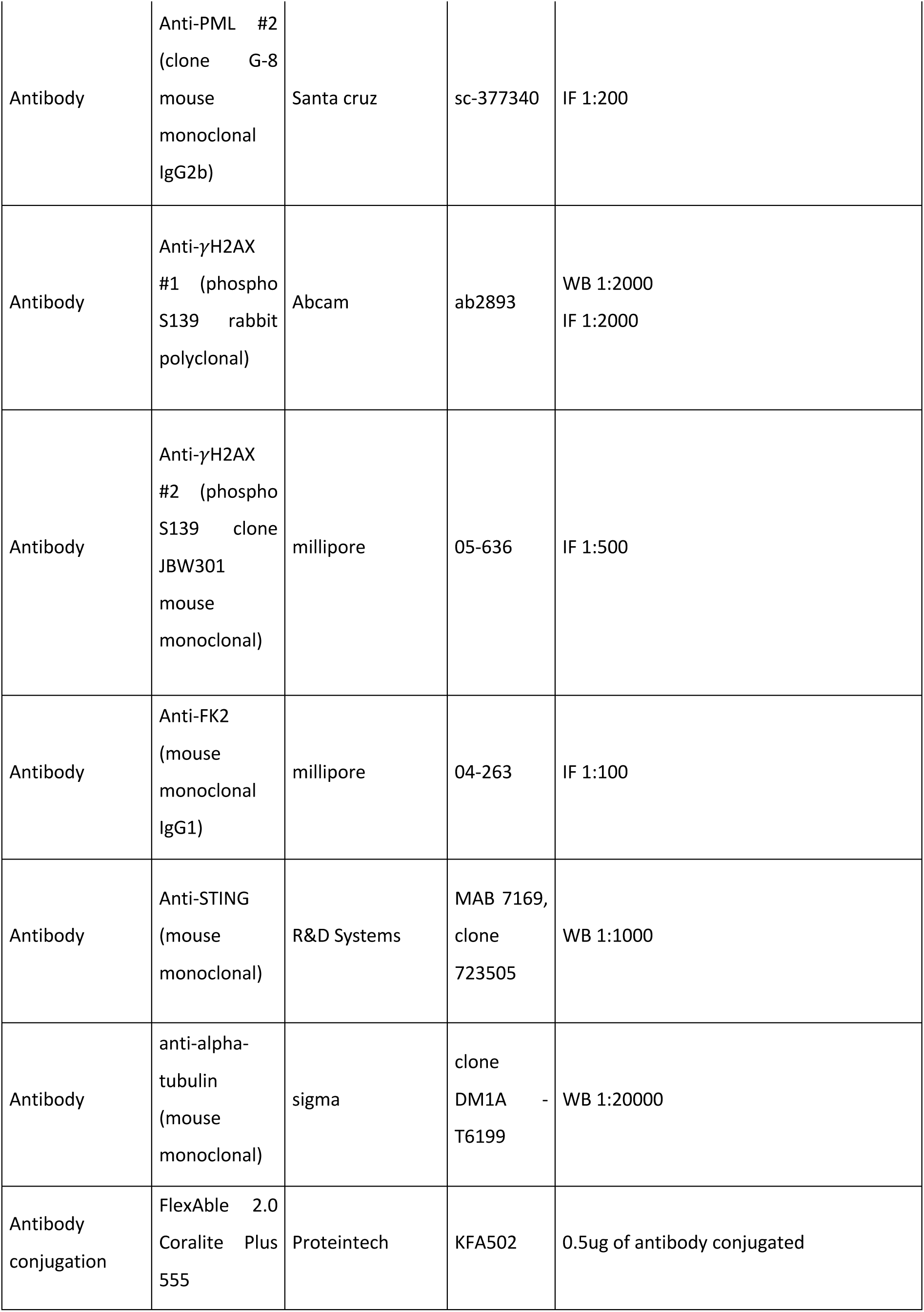
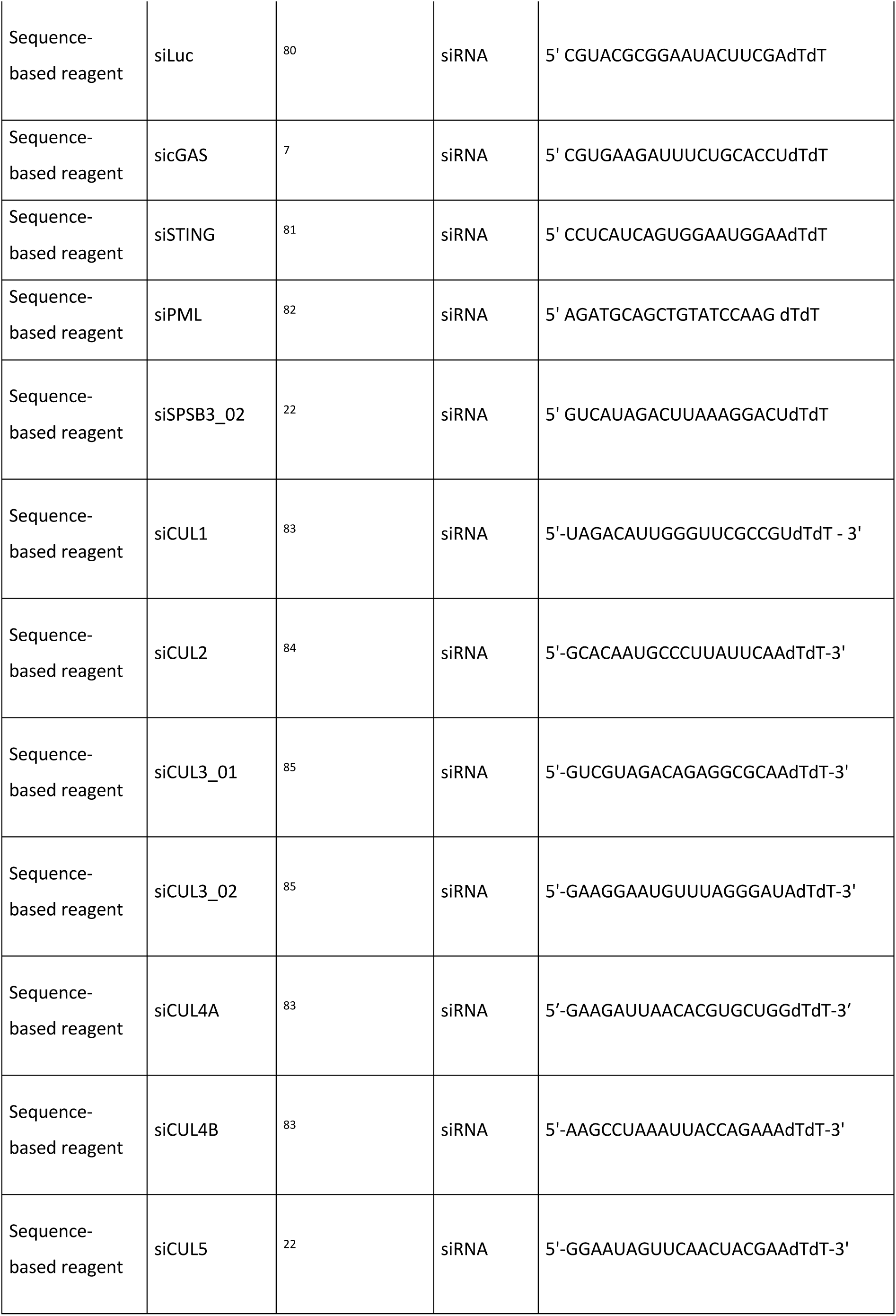
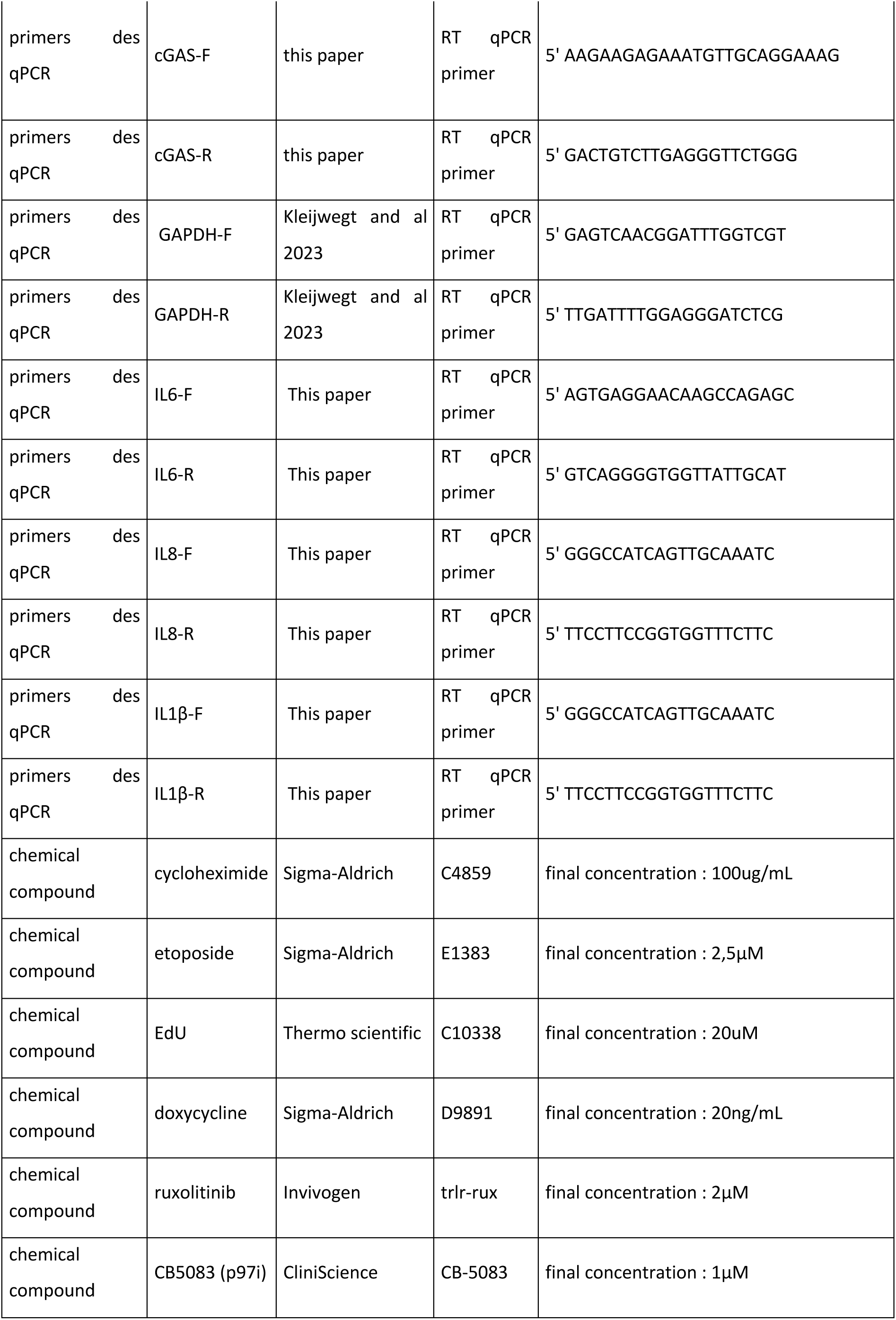
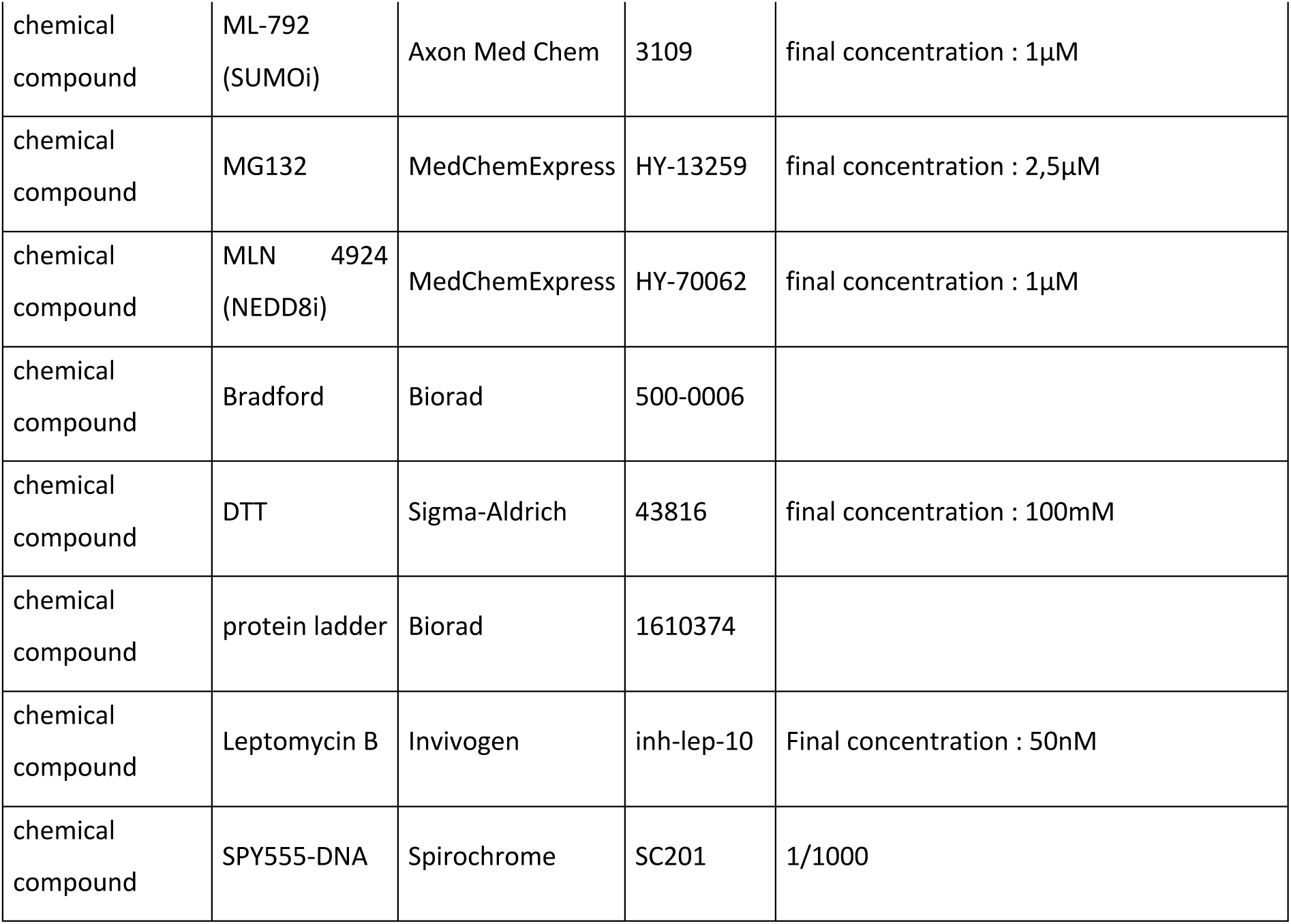

## Acknowledgments

PL laboratory is funded by grants from the Centre National de la Recherche Scientifique (CNRS), Institut National de la Santé et de la Recherche Médicale (INSERM), University Claude Bernard Lyon 1, French National Agency for Research-ANR [EPIPRO ANR-18-CE15-0014-01, CHROMACoV ANR-20-COV9-0004; IFN-Epi-IM ANR-21-CE17-0018]; LabEX

DEVweCAN/DEV2CAN [ANR-10-LABX-61]; AFM-Téléthon Plans stratégiques MyoNeurALP & MyoNeurALP2, CNRS Prematuration program 2021-2023, the Comité départemental du Rhône de La Ligue contre le Cancer and l’Association pour la Recherche contre le Cancer (4th year PhD grant to FB). PL is a CNRS Research Director and AC is associate professor in the University Claude Bernard Lyon 1. AC is also a junior member of the Institut Universitaire de France, and funded by this organization. The authors also greatly acknowledge Bruker for collaboration with CNRS.

## Author contributions

A.C. and P.L. designed the study, directed its implementation, analyzed and interpreted the data, and wrote the manuscript. F.B., T.B., and C.K. performed experiments, analyzed the data and reviewed the manuscript. A.C. performed the CUT&RUN experiments and data analysis.

L.B. and K.M. performed the dSTORM image acquisition and analysis and reviewed the manuscript. W.B. analyzed the RNA-Seq data and reviewed the manuscript.

## Competing interests

The authors declare no competing interests.

**Supplementary Figure 1:** (A) Histograms shows relative *cGAS* mRNA levels (normalized on β*-actin* mRNA levels) of BJ cells treated or not with etoposide at 10uM for 24h. Rationalization was performed on the non-treated condition. P-values (Student t-test) : * <0,05. Numbers represent the mean of three independent experiments. (B) Fluorescence microscopy visualization of cGAS (red) in BJ cells treated with etoposide at 10μM for 24h and leptomycinB at 50nM for 24h. Cell nuclei are visualized by DAPI staining (grey). Scale bar is 10μm. (C) (Left) Western-blot visualization of cGAS and GFP-cGAS from total cell extracts on BJ GFP-cGAS cells treated with Doxycycline at 20ng/mL for 24h. Actin is a loading control. (Right) Fluorescence microscopy visualization of GFP-cGAS (green) in cells treated as on the left panel. Cell nuclei are visualized by DAPI staining (grey). Scale bar is 10μm. (D)(Left) Fluorescence microscopy visualization of cGAS (red) in HeLa cells treated or not with etoposide at 10μM for 24h. Cell nuclei are visualized by DAPI staining (grey). Scale bar is 10μm. (Right) Dot plot shows quantitative analysis of nuclear cGAS intensity in cells. P-values (Student t-test) : ****<0,0001. (E) Western-blot visualization of cGAS from total cell extracts on HeLa cells treated as in (D). Actin is a loading control and quantification of cGAS levels relative to actin are shown below the WB (numbers are representative from three independent experiments). (F) Western-blot visualization of cGAS from different fractions of HeLa cells treated with etoposide at 10μM for 24h. Actin is a loading control and quantification of cGAS levels relative to actin are shown below the WB.

**Supplementary Figure 2:** (A) Western-blot visualization of cGAS and γH2AX from total cell extracts on BJ cells treated with TSA at 2μM for 24h and MG132 at 2,5μM for 8h. Actin is a loading control and quantification of cGAS and γH2AX levels relative to actin are shown below the WB. (B) Fluorescence microscopy visualization of PML (green) and cGAS (red) in BJ cells treated as in (A). Cell nuclei are visualized by DAPI staining (grey). Scale bar is 10μm. Insets represent enlarged images (3X) of selected areas. (C) Fluorescence microscopy visualization of ACA (green) and cGAS (red) in BJ cells treated with etoposide at 10μM for 24h and with MG132 at 2,5μM for 8h. Cell nuclei are visualized by DAPI staining (grey). Scale bar is 10μm. (D) Fluorescence microscopy visualization of H2A-H2B-GFP (green) using an H2A-H2B Nanobody fused to EGFP and DNA (red) stained with SPY-DNA-555 in untreated BJ cells. Scale bar represents 10μm. (E) Fluorescence microscopy visualization of PML (green), cGAS (red) and γH2AX (cyan) in BJ cells treated with etoposide at 10μM for 24h and with MG132 at 2,5μM for 8h. Cell nuclei are visualized by DAPI staining (grey). Scale bar represents 10μm. Insets represent enlarged images (3X) of selected areas.

**Supplementary Figure 3:** (A) Fluorescence microscopy visualization of FK2 (green) and PML (red) in BJ cells treated with etoposide at 10μM for 24h and with MG132 at 2,5μM for 8h. Cell nuclei are visualized by DAPI staining (grey). Scale bar represents 10μm. Insets represent enlarged images (3X) of selected areas. (B) Fluorescence microscopy visualization of FK2 (green) and cGAS (red) in BJ cells treated as in (A). Cell nuclei are visualized by DAPI staining (grey). Scale bar represents 10μm. Insets represent enlarged images (3X) of selected areas. (C) Fluorescence microscopy visualization of cGAS (green), PML (cyan) and ADRM1 (red) in non treated BJ cells. Insets represent enlarged images (3X) of selected areas. Scale bar is 10μm.

**Supplementary Figure 4:** (A) Histograms show RT-QPCR analysis of mRNA levels for the indicated genes in BJ cells treated with the indicated siRNAs for 48h (n=2 to 3 independent experiments). P-values (Student t-test) : ****<0,0001. (B) Histogram shows the percentage of cells with cGAS foci juxtaposing PML NBs (n=3 independent experiments). P-values (Student t-test) : *<0,05; ****<0,0001 ; ns : non-significant. (C) Fluorescence microscopy visualization of PML (green) and cGAS (red) in BJ cells treated the indicated siRNAs for 48h. Cell nuclei are visualized by DAPI staining (grey). Scale bar represents 10μm. (D) (Left) Fluorescence microscopy visualization of PML (green) and cGAS (red) in BJ cells treated with etoposide at 10μM for 24h and with MG132 at 2,5μM for 8h and with the indicated siRNAs for 48h. Cell nuclei are visualized by DAPI staining (grey). Scale bar represents 10μm. Insets represent enlarged images (3X) of selected areas. (Right) Histogram shows quantitative analysis of cells with cGAS foci from 2 independent experiments. (E) Western-blot visualization of Cullin3 and Actin from total cell extracts on BJ cells treated with etoposide at 10μM for 24h and with the indicated siRNAs for 48h. Actin is a loading control. (F) Fluorescence microscopy visualization of PML (green) and cGAS (red) in BJ cells treated the indicated siRNAs for 48h. Cell nuclei are visualized by DAPI staining (grey). Scale bar represents 10μm.

**Supplementary Figure 5:** (A) Fluorescence microscopy visualization of PML (green) and cGAS (red) in BJ cells treated with p97i (CB5083) at 1μM for 24h. Cell nuclei are visualized by DAPI staining (grey). Scale bar represents 10μm. Insets represent enlarged images (3X) of selected areas. (B) Western-blot visualization of cGAS and Actin from total cell extracts on BJ cells treated with TSA at 2μM for 24h, and p97i (CB5083) at 1μM for 24h. Actin is a loading control and quantification of cGAS levels relative to actin are shown below the WB. (C) Fluorescence microscopy visualization of cGAS (red) in BJ cells treated as in (B). Cell nuclei are visualized by DAPI staining (grey). Scale bar represents 10μm. (D) Western-blot visualization of cGAS and H3 in the chromatin-bound fraction of BJ cells treated as in (B). Quantification of cGAS levels relative to H3 are shown below.

**Supplementary Figure 6:** (A) CUT&RUN against IgG, H3K4me3 or endogenous cGAS was performed in BJ primary cells, left untreated (IgG, H3K4me3, cGAS_NT), treated with etoposide at 10μM for 24h (cGAS_eto) or treated with etoposide at 10μM for 24h and MG132 at 2,5μM for the last 8h (cGAS_eto_MG). Plot Profile shows density of the indicated conditions across human promoters. (B) Plot Profile shows density of the indicated conditions on identified H3K4me3 peaks using MACS2. (C) Genome browser snapshot of the indicated conditions across the indicated region. (D) Genome browser snapshot of the indicated conditions showing no enrichment of cGAS across standard genes of chromosome 19. (E) Plot Profile shows density of cGAS in the indicated conditions on KZFP genes. While cGAS enrichment diminished on KZFPs with the eto_MG condition, arrows indicate spreading of the signal beyond KZFPs. (F) Plot Profile shows density of cGAS in the indicated conditions on ERVL or ERV1.

**Supplementary Figure 7:** (A) Histogram shows quantitative analysis of cells with EdU incorporation on BJ cells treated or not with etoposide at 10μM for 24h and recovered for 5 days, 7 days or 10 days. (B) Histograms show relative *IL-6*, *IL-8* or *IL1-*β mRNA levels (normalized on β*-actin* mRNA levels) of BJ cells treated or not with etoposide at 10μM for 24h and recovered for 7 days or 10 days. Rationalization was performed on the non treated condition. (C) Bright-field microscopy visualization of SA-βgal on BJ cells treated with etoposide at 10μM for 24h and recovered for 10 days. (D) Histogram shows quantitative analysis of cells with HIRA localization at PML NBs on BJ cells treated with etoposide at 10μM for 24h and recovered for 5 days, 7 days or 10 days. (E) (Left) Fluorescence microscopy visualization of PML (green) and cGAS (red) in BJ cells treated with NCS at 200ng/mL for 20min and recovered for 1h. Cell nuclei are visualized by DAPI staining (grey). Scale bar is 10μm. Insets represent enlarged images (3X) of selected areas. (Right) Histogram shows quantiative analysis of cells with HIRA localization at PML NBs. (F) (Left) RNA-Seq data from BJ cells treated with etoposide for 24h, 72h, or left untreated (NT). Heatmap of the expression of differentially expressed genes between NT and eto 24h in the different conditions. (Right) Bubble chart showing KEGG pathway enrichment analysis of differentially expressed genes (DEGs) upregulated or downregulated in the eto 24h condition compared to the NT condition. (G) Western-blot visualization of cGAS, STING, Tubulin and Actin from total cell extracts on BJ cells treated with etoposide at 10μM for 24h and with the indicated siRNAs for 48h. Tubulin and Actin are loading controls and quantification of cGAS or STING levels relative to actin are shown below the WB (numbers are representative from three independent experiments). (H) Fluorescence microscopy visualization of HIRA (green), and PML (red) in BJ cells treated with etoposide at 10μM for 24h and with the indicated siRNAs. Cell nuclei are visualized by DAPI staining (grey). Scale bar is 10μm. Insets represent enlarged images (3X) of selected areas. (I) Histogram shows quantitative analysis of cells with HIRA localization at PML NBs (n=3 independent experiments) in cells treated as in (H). P-values (Student t-test) : ***<0,001, ns: non significant.

**Supplementary Figure 8 :** (A) Fluorescence microscopy visualization of γH2A.X (green) and cGAS (red) in BJ cells treated with etoposide at 10μM for 24h and with p97i (CB5083) at 1μM for 24h. Cell nuclei are visualized by DAPI staining (grey). Scale bar represents 10μm.

