## Supplementary figures and images for "A p97-PML NBs axis is essential for chromatin-bound cGAS untethering and degradation upon senescence-prone DNA damage"

### Supplementary Figures

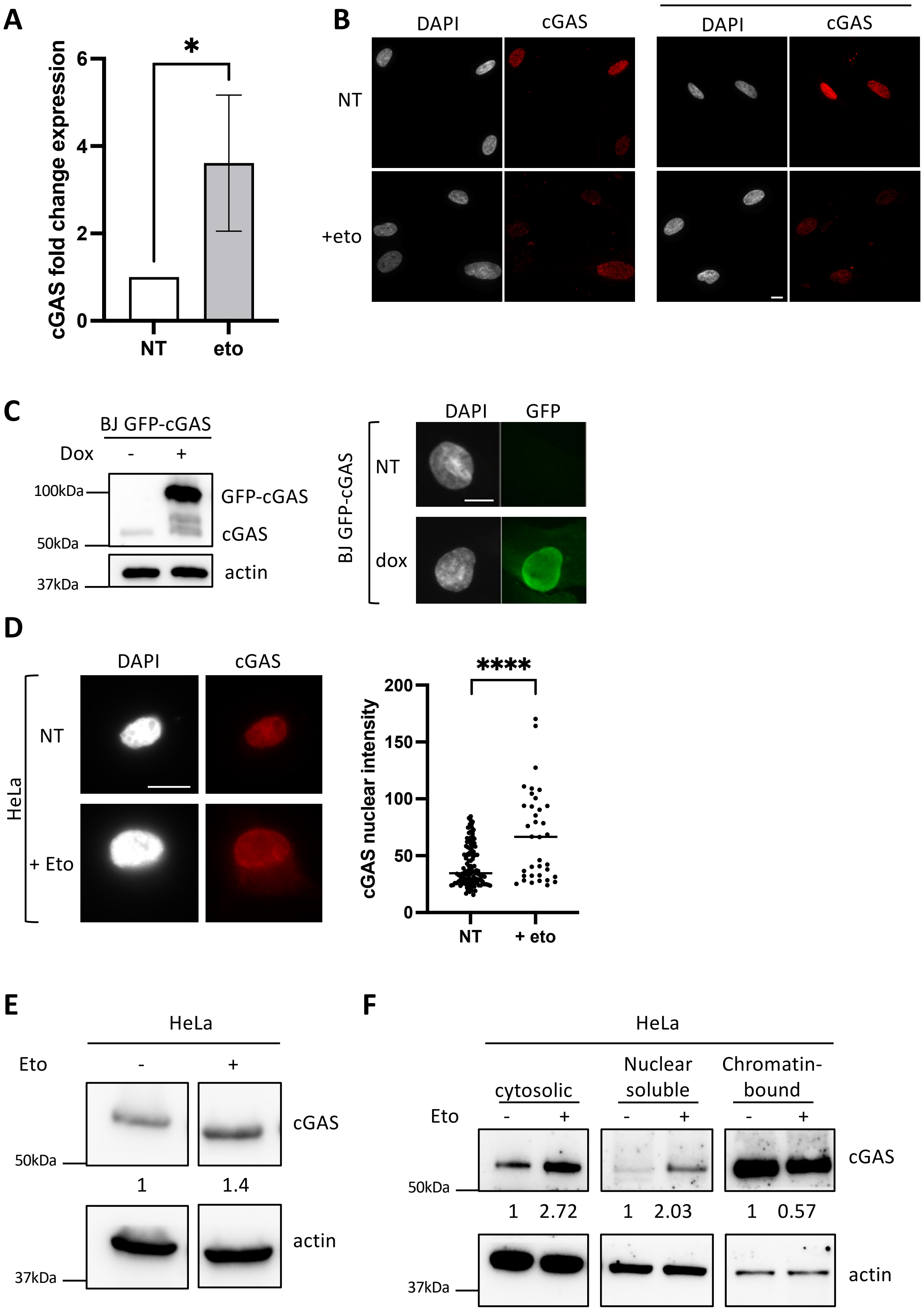

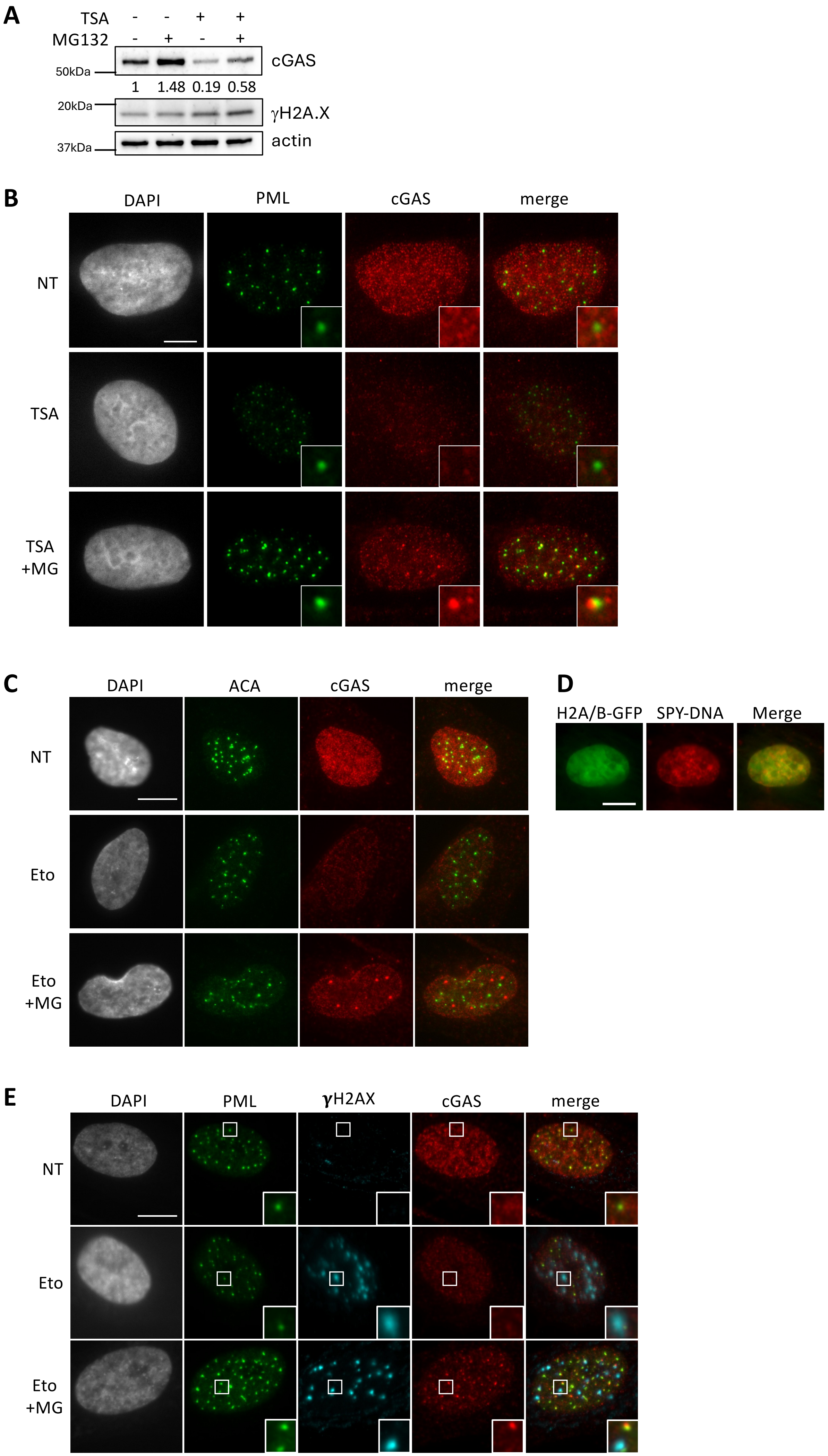

#### Sup. Figure 3

A

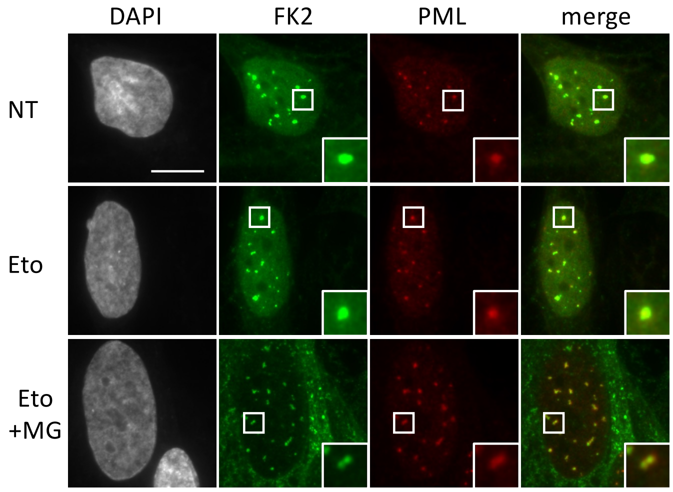


B

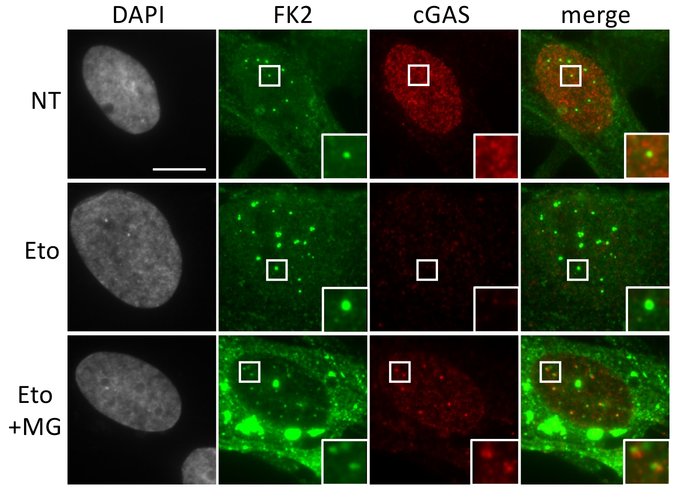


C

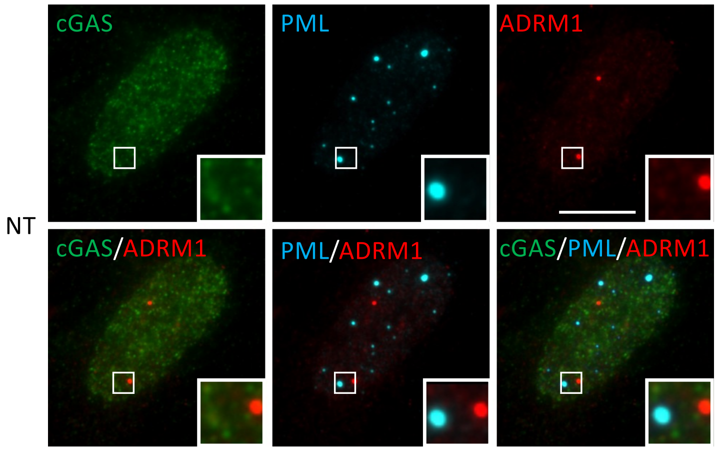

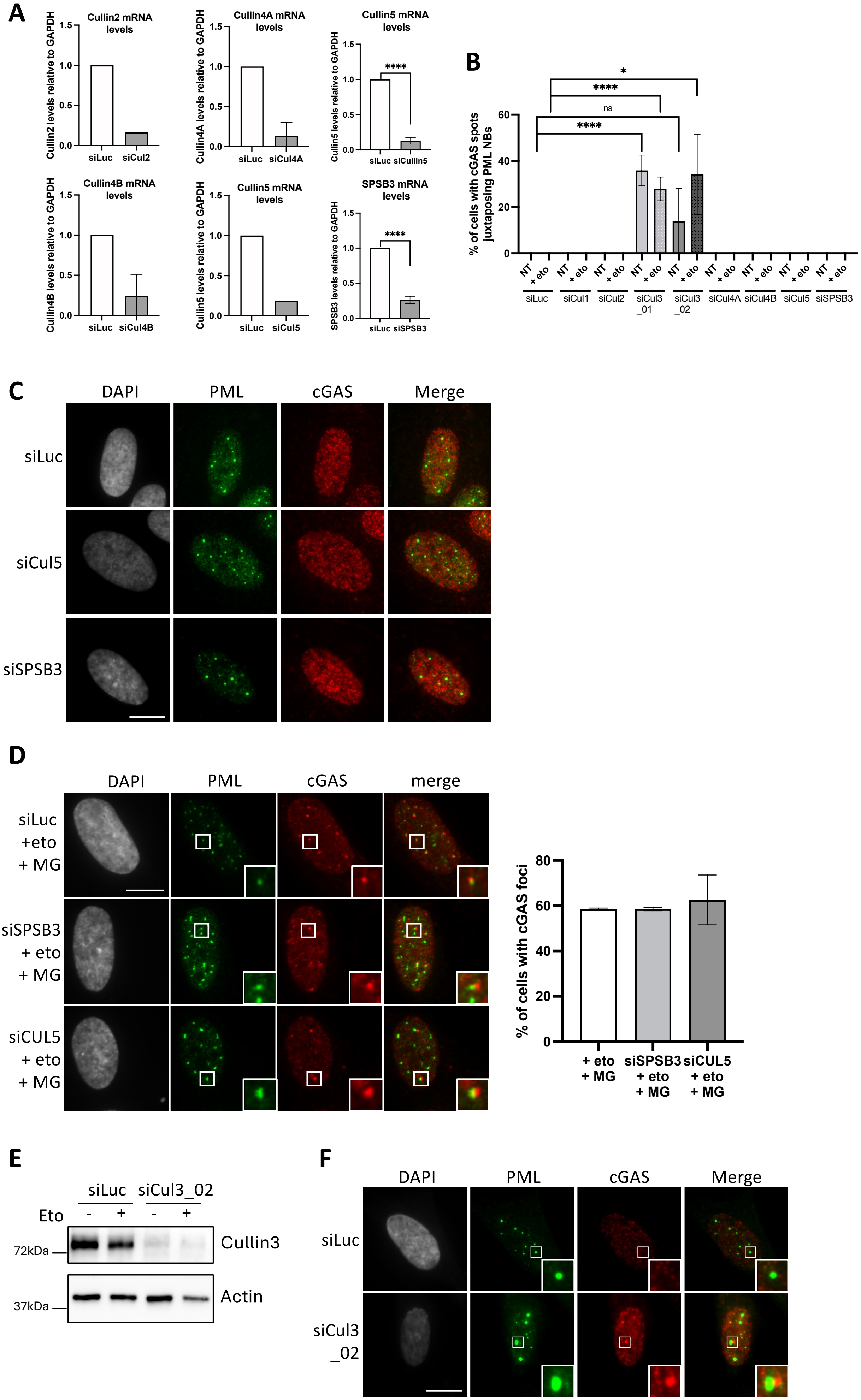

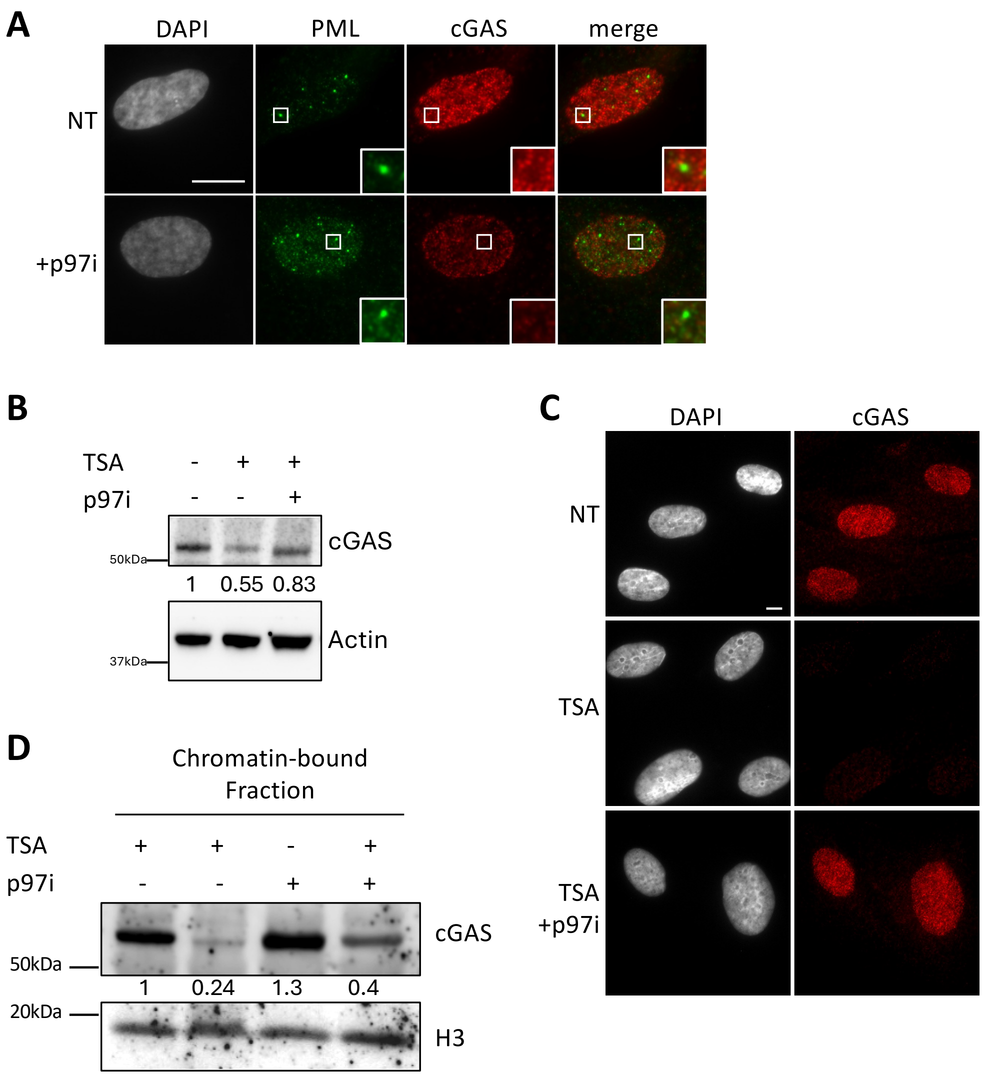

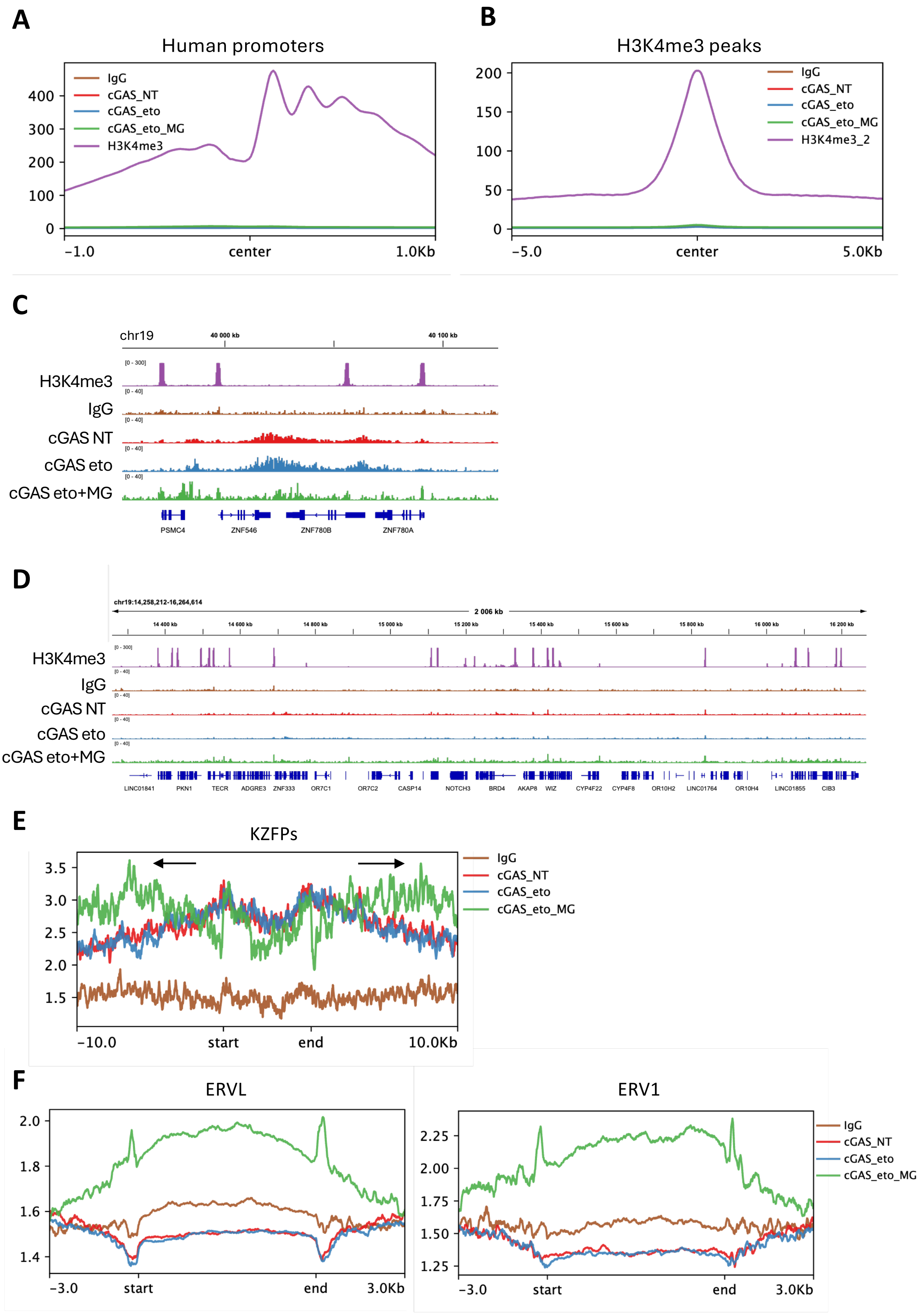

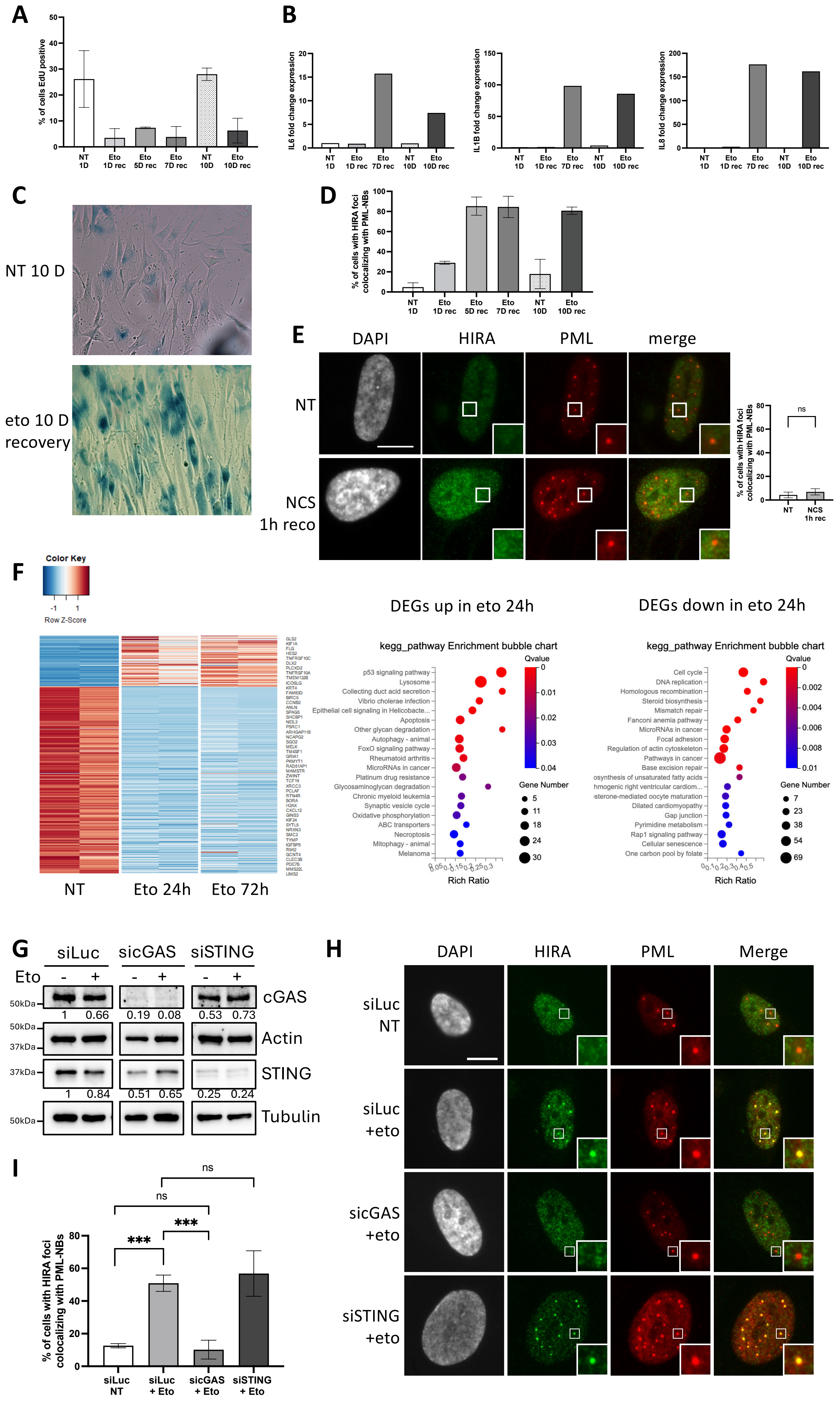

**Sup. Figure 8**

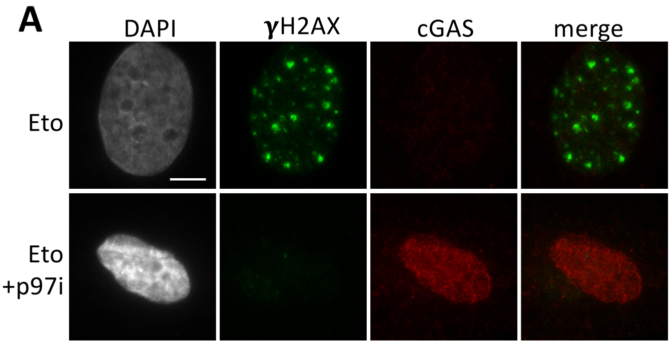
